# Basal forebrain cholinergic neurons are part of the threat memory engram

**DOI:** 10.1101/2021.05.02.442364

**Authors:** Prithviraj Rajebhosale, Mala Ananth, Richard Crouse, Li Jiang, Gretchen López- Hernández, Christian Arty, Shaohua Wang, Alice Jone, Chongbo Zhong, Niraj S. Desai, Yulong Li, Marina R. Picciotto, Lorna W. Role, David A. Talmage

## Abstract

Although the engagement of cholinergic signaling in threat memory is well established (Knox, 2016a), our finding that specific cholinergic neurons are requisite partners in a threat memory engram is likely to surprise many. Neurons of the basal forebrain nucleus basalis and substantia innominata (NBM/SI_p_) comprise the major source of cholinergic input to the basolateral amygdala (BLA), whose activation are required for both the acquisition and retrieval of cued threat memory and innate threat response behavior. The retrieval of threat memory by the presentation of the conditioning tone alone elicits acetylcholine (ACh) release in the BLA and the BLA-projecting cholinergic neurons manifest immediate early gene responses and display increased intrinsic excitability for 2-5 hours following the cue-elicited memory response to the conditioned stimulus. Silencing cue-associated engram-enrolled cholinergic neurons prevents the expression of the defensive response and the subset of cholinergic neurons activated by cue is distinct from those engaged by innate threat. Taken together we find that distinct populations of cholinergic neurons are recruited to signal distinct aversive stimuli via the BLA, demonstrating exquisite, functionally refined organization of specific types of memory within the cholinergic basal forebrain.

## Introduction

Acetylcholine (ACh) is critical for cognition. The basal forebrain cholinergic system (BFCS), the major source of acetylcholine to the cortex, hippocampus, and amygdala, is composed of few cholinergic neurons which are sparsely distributed along the base of the forebrain. These neurons send long projections that are extensively branched in their target fields (Zaborszky et al., 2012). Given this, relatively few cholinergic neurons must be both anatomically and functionally organized in order to modulate large, behaviorally relevant circuits (Gielow and Zaborszky, 2017; Zaborszky et al., 2015).

ACh in the central nervous system acts as a modulator of ongoing synaptic transmission. Early impressions were that cholinergic transmission was slow, working at timescales of hundreds of milliseconds to seconds, providing broad control over brain states such as arousal, sleep, and attention ((Ballinger et al., 2016; Picciotto et al., 2012). Technological advances have refined our understanding, revealing that ACh can signal over a range of timescales, providing mechanisms for both fast and slow signaling, determined by several factors including location within the brain, mechanism of release, and type of synaptic contact (Disney and Higley, 2020). Disruptions to this normal cholinergic transmission are thought to contribute to a number of neuropsychiatric disorders (Higley and Picciotto, 2014; Sarter et al., 1999) and lead to altered behavior in rodents (Crouse et al., 2020; Hersman et al., 2017; Jiang et al., 2016).

Acetylcholine plays an important role in modulating emotionally salient memories (Ballinger et al., 2016; Knox, 2016b; Luchicchi et al., 2014). We and others have found that cholinergic signaling in the basolateral amygdala (BLA) is important for generating defensive behaviors in response to both learned and innate threats (Jiang et al., 2016; Power and McGaugh, 2002; Wilson and Fadel, 2017). Optogenetic manipulation of endogenous ACh release in the BLA during learning modulates the expression of threat response behaviors in mice upon recall of a conditioned stimulus (Jiang et al., 2016). Stimulating release of ACh increased activity of BLA principal neurons, in part by increasing the release probability of glutamatergic inputs to these neurons, and was sufficient to induce long-term potentiation (LTP) when paired with minimal (non-LTP generating) stimulation of glutamatergic input to the BLA (Jiang et al., 2016; Unal et al., 2015). Interestingly, BLA principal neurons that become activated during threat learning and recall (as marked by immediate-early gene (IEG)-based markers), also display an increase in the probability of presynaptic glutamatergic transmission (Nonaka et al., 2014). Thus, memory formation and retrieval are associated with fast synaptic mechanisms that are modulated by ACh, which in turn is necessary for the proper learning and expression of threat response behaviors. Given the broad distribution of cholinergic input across the BLA, and the well-established role of ACh in modulating BLA plasticity, the basal forebrain cholinergic system is well-positioned to serve an important role in the encoding of threat memories and generation of threat response behaviors.

Neuronal ensembles in the amygdala are activated during threat learning. Recall of threat memory is associated with reactivation of those same ensembles, thereby earning them the title of “engram cells”. Additional studies have found that activated neurons display increased excitability, potentially improving the chance that these neurons will enroll into the engram (DeNardo and Luo, 2017; Josselyn et al., 2015; Liu et al., 2012; Reijmers et al., 2007; Tonegawa et al., 2015; Yiu et al., 2014). These changes in excitability have been attributed to a number of mechanisms including spine dynamics, changes in intrinsic excitability and synaptic plasticity (Barth et al., 2004; Butler et al., 2018; Kim et al., 2014; Liu et al., 2012; Nonaka et al., 2014; Pignatelli et al., 2019). Of these, synaptic plasticity in BLA neurons has been causally linked to associative learning and memory recall *in vivo* (Nabavi et al., 2014). Taken together with findings on cholinergic function in the BLA, these findings raise the question of how and when cholinergic neurons are recruited on behaviorally relevant timescales for associative learning.

In this study we asked how distinct populations of basal forebrain cholinergic neurons contribute to threat responses. Using a genetically encoded ACh sensor, genetic tagging, chemogenetic manipulations and electrophysiological recordings, we identified a population of BLA-projecting BFCNs that are required for learned threat responsiveness and the establishment of a threat memory engram. We also found that distinct populations of BFCNs are necessary for freezing behavior following cue conditioned threat learning than for freezing behavior elicited by an innately threatening stimulus. Based on our findings, we propose that NBM/SI_p_ BFCNs are an essential part of the threat memory engram and provide evidence for previously unrecognized functional organization in the BFCS.

## Results

### Acetylcholine is released in the basal lateral amygdala in response to threat

Animals recognize varied sensory stimuli and categorize them as either threatening or non-threatening. Recognition of threatening stimuli can be innate, or acquired, for example, by association of an aversive experience with an innocuous, co-occurring sensory input. In this study we sought to understand if, and when, the basal forebrain cholinergic system is engaged during associative threat learning and recall and in response to innate threat.

The first question we asked was whether acetylcholine was released in the BLA during associative threat learning (popularly known as fear conditioning or Pavlovian conditioning). We focused these studies on the BLA because of the well-established role this region plays in learning, and in the generation of threat responses. We have previously demonstrated that silencing cholinergic input to the BLA during cue-conditioned threat learning (pairings of naïve tone with foot shock) blunts subsequent defensive responses to the conditioned stimulus (tone) (Jiang et al., 2016).

To monitor acute changes in extracellular ACh levels during the cue conditioned threat learning task, we expressed a genetically encoded ACh sensor, GRAB_ACh3.0_ in BLA neurons and visualized fluorescence using fiber photometry (**Figure 1A left**) (Jing et al., 2020; Jing et al., 2018). Our associative threat learning protocol (**Figure 1A right**) involved placing animals in a novel chamber and exposing them to an 80dB tone for 30 sec. During the final 2 sec of the tone the animals received a foot-shock (0.7 mA). The tone-shock pairing was repeated twice (for a total of 3 pairings). Twenty-four hours later, mice were placed in a different chamber and exposed to tone alone.

**Figure 1:**
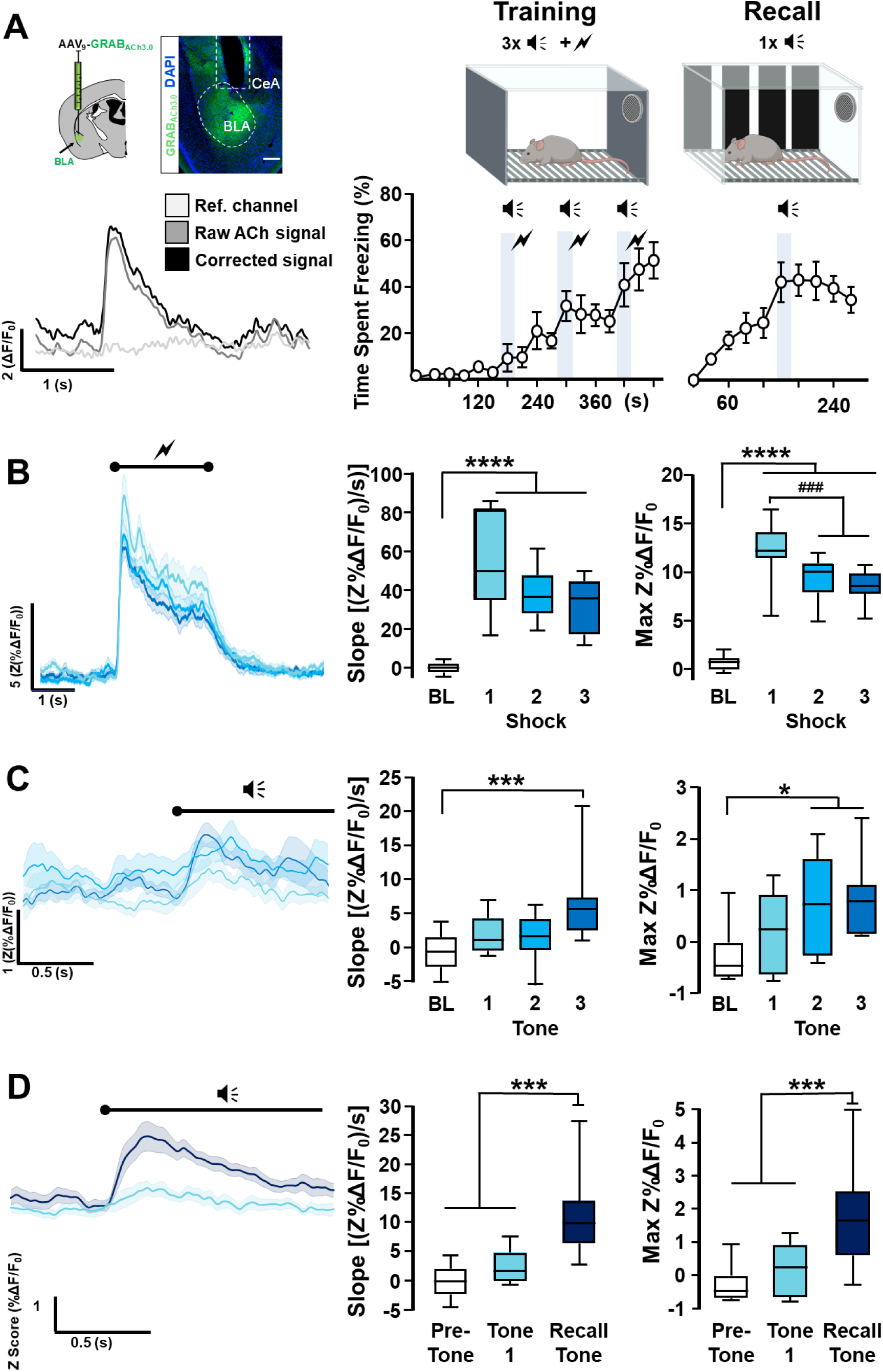
Acetylcholine is released in BLA during training and during threat recall (see also Figure S1). **A. Left.** Strategy for targeting the genetically encoded ACh sensor (GRAB_ACh3.0_) to BLA. Image of ACh sensor expression (green). White dotted oval delineates ACh sensor labeled BLA. White dotted box denotes prior location of optical fiber. Scale bar = 100µm. Recording of ACh release in response to a single, 0.5s footshock. Traces depict corrected (black) and uncorrected (gray) ΔF/F_0_ signal compared to the reference channel (light gray). **Right.** Schematic of the associative threat learning protocol consisting of 3 tone + shock pairings during the training period, and tone alone during the recall session. Quantification of freezing behavior throughout the training session (30s time-bins; n=9) and during the recall session (30s time-bins; n=7). **B.** Average traces of ACh release in response to 2 sec foot <u>shock</u> (∼) during tone-shock training (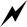); shading represents SEM. Quantification of change in slope and maximum ACh release during baseline (BL) period (white box), and in response to each shock (Tone1: turquoise blue, Tone2: deep sky blue, Tone3: navy blue). The response to all 3 shocks differed from baseline (slope (Z%ΔF/F0)/s) and max (Max Z%ΔF/F0) RM one-way ANOVA ****, p<0.0001). The maximum release in response to the 1^st^ shock differed from the response to shocks 2&3 (###, p<0.005), shock 1 v. 2 (p=0.0032) and shock 1 v. 3 (p=0.0006). p-values were corrected for multiple comparisons using Tukey’s post-hoc correction. Power=1. **C.** ACh release in response to <u>tone</u> (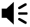) during training (n=11). Average traces of ACh release in response to tone; shading represents SEM. Quantification of change in slope and maximum ACh release during baseline period (white box) and in response to onset of the tone (Tone1: turquoise blue, Tone2: deep sky blue, Tone3: navy blue) for each tone + shock pairing. The change in slope in response to the third tone differed significantly from baseline (Friedman test, Dunn’s corrected***, p<0.005). Power=0.96. The peak of release in response to tones 2 / 3 differed from baseline (Friedman test, Dunn’s corrected *, p<0.05). Power=0.97. **D.** ACh release in response to conditioned threat recall (n= 11). Average traces of ACh release in response to tone; shading represents SEM: naïve tone (tone 1 during training) in turquoise blue, recall tone in Prussian blue (onset of tone set to T=0; n=11). Quantification of change in slope and maximum ACh release during baseline period, in response to the first (naïve) tone and in response to the recall tone. The response to the recall tone was significantly greater than to the pre-tone period or naïve tone (tone 1) (slope: ****, p<0.0001; peak: **** p<0.0001) (slope & peak: Baseline v. Tone1, p=0.859; Baseline v. Recall, p=0.0002; Tone1 v. Recall, p=0.0085). Friedman test, Dunn’s corrected p-values. Power= 0.99. *See also Figure S1.

Acute foot shock increased ACh release in the BLA (**Figure 1B**). The slope of the rise in GRAB_ACh3.0_ fluorescence (**Figure 1B**) was not significantly different between the three foot-shocks (p=0.105) but was significantly higher compared to baseline (p<0.0001). The magnitude of changes in GRAB_ACh3.0_ fluorescence in response to each of the three foot shocks (**Figure 1B right**) was highest for the first shock (p<0.0001) with a significant reduction in the peak amplitude for the second (Shock 1 vs. Shock 2 p=0.0032; Shock 2 vs. Baseline p<0.0001) and third shocks (Shock 1 vs. Shock 3 p=0.0006, Shock 2 vs. Shock 3 p=0.3103; Shock 3 vs. Baseline p<0.0001).

When we examined the response to the three tones during the training session (**Figure 1C**) two results stood out. First, there was no significant response to naïve tone (i.e. to the tone before pairing with footshock) (Slope: baseline v. Tone 1 p>0.9999: Peak: baseline v. Tone 1 p>0.9999), or to tone 2 (Slope: baseline v. Tone 2 p=0.5919; Peak: baseline v. Tone 2, p=0.0494). Second, tone 3 was associated with both a significant increase in the slope from baseline to peak response (**Figure 1C middle**, baseline v. Tone 3 p=0.0017) and in peak fluorescence (**Figure 1C right**, baseline v. Tone 3 p=0.0494).

Twenty-four hours after tone-shock pairing, we measured GRAB_ACh3.0_ fluorescence in response to the tone alone in a novel context (recall session; **Figure 1D**). The animals now showed a significant increase in the slope (**Figure 1D**; Baseline v. Recall p=0.0002; Tone 1 v. Recall p=0.0085) and in the overall magnitude of ACh release (**Figure 1D**; Baseline v. Recall p=0.0002; Tone 1 v. Recall p=0.0085) in response to tone. The change in tone-associated ACh release required pairing with foot shock: none of three consecutive tones alone (without shock), or a subsequent tone presentation after 24 hr, induced significant changes in ACh release in the BLA (**Figure S1A&S1B**).

### NBM / SI cholinergic neurons are activated by threat learning and reactivated during threat memory recall

Following associative threat learning, BLA-projecting cholinergic neurons showed changes in ACh release in the BLA in response to a previously innocuous auditory stimulus; this response occured exclusively following pairing with a shock. These results led us to ask whether a specific population of cholinergic neurons in the basal forebrain participate in the threat memory engram. We began our investigation in the NBM/SI_p_ given its established input to the BLA (Zaborszky et al., 2012).

To investigate whether NBM/SI_p_ cholinergic neurons are part of a threat memory engram, we asked whether there was a population of cholinergic neurons activated during training that were reactivated during recall. We injected the offspring of a cross of Chat-IRES-Cre x Fos-tTA:Fos-shGFP with AAV_9_-TRE-DIO-mCherry-P2A-tTA^H100Y^ (resulting in activity (tTA) dependent, Cre dependent (ADCD) mCherry expression, see methods and **Figure S2** for virus details). These mice carry three transgenes – Cre recombinase expressed in cholinergic neurons, the doxycycline (Dox) repressible, tetracycline transactivator (tTA) expressed following activation of the *fos* promoter, and a destabilized green fluorescent protein (short half-life GFP), also under transcriptional regulation of the *fos* promoter. tTA and shGFP are transiently expressed in activated neurons. In the absence of Dox, activation of Cre-expressing cholinergic neurons leads to the continuous expression of the virally transduced mCherry.

The mice were assigned to one of the following groups: 1. home cage throughout, 2. exposed to tone without foot shock (tone alone), or 3. standard threat learning paradigm (tone+shock). Twenty-four hours prior to the training session they were switched from Dox-containing to Dox-free chow to allow function of tTA, and then immediately placed back on Dox-containing chow after the tone-shock pairings (**Figure 2A**). This switch from Dox on-off-on was also performed for mice that remained in their home cages and for those that were trained without shock.

**Figure 2.**
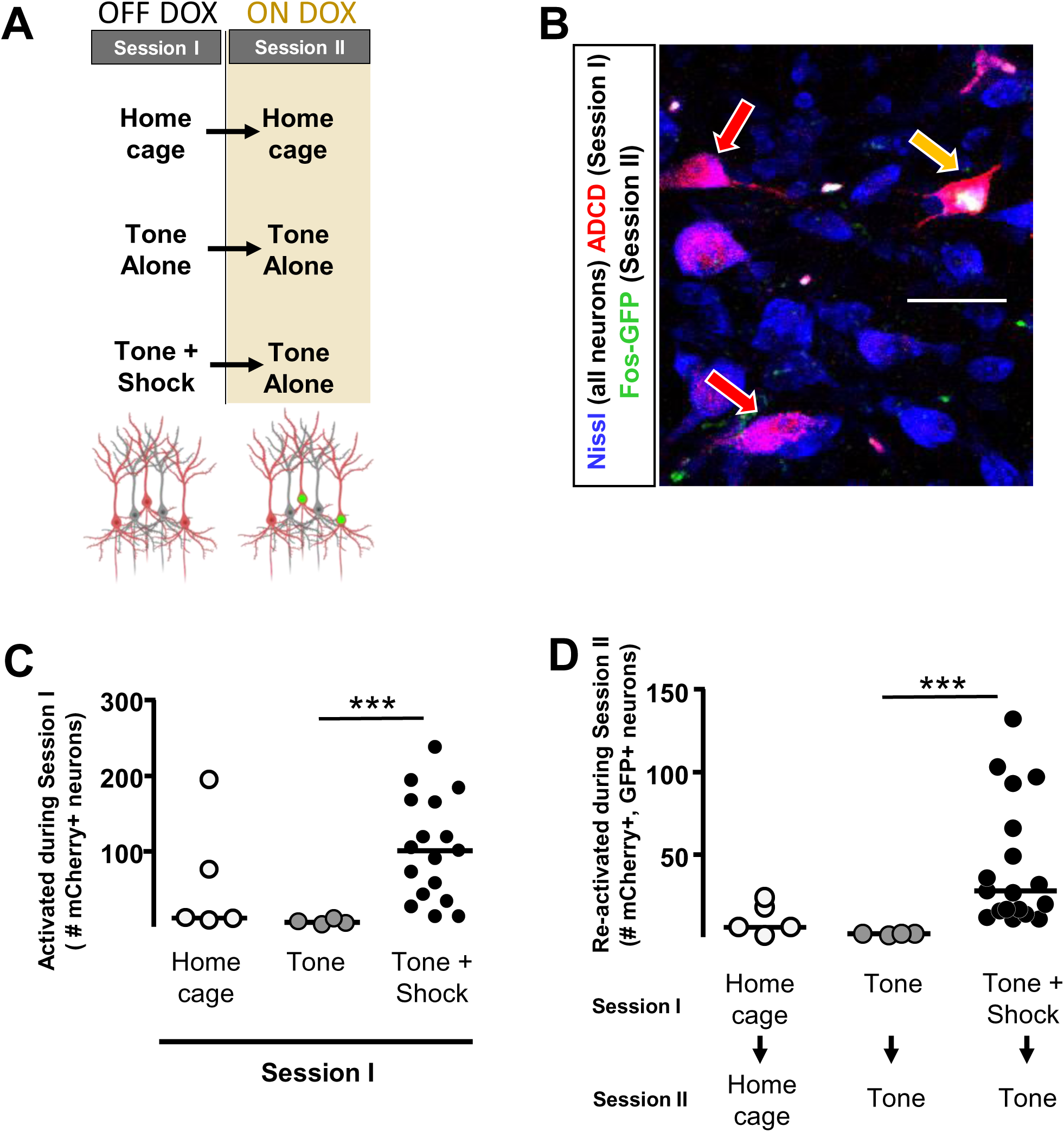
Activity-dependent labeling of NBM/SI cholinergic neurons (see also Figure S2) **A.** Strategy for labeling activated NBM/SI_p_ cholinergic during training <u>and</u> recall. Animals were injected in the NBM/SI_p_ with ADCD-mCherry virus. During Session 1 (off Dox) animals either remained in their home cage, were exposed to 3 tones (Tone alone), or were exposed to 3 tone-shock pairings. During session 2 (recall session), animals remained in home cage or were exposed to a single tone. Cholinergic neurons activated during training express mCherry (red during training), and cells activated during recall transiently express GFP (green during recall). **B.** Image of the NBM/SI_p_ showing cholinergic neurons activated during training (**red arrow**) or both training and recall (reactivated – **yellow arrow**; image taken at A/P ∼ −0.8 from Bregma; scale bar 25 μm). **C.** Quantification of cholinergic neurons activated during Session 1: home cage (n = 5 sections from 4 mice), tone alone (n=4 sections from 2 mice), and tone + shock (n= 17 sections from 8 mice). Significantly more cholinergic neurons expressed mCherry following tone-shock pairings (Kruskal-Wallis p<0.01). Tone-shock compared to tone only (***, p=0.004, Tukey corrected). **D.** Quantification of **reactivated** cholinergic neurons (activated both during Session 1 (training) <u>and</u> during Session 2 (recall)). Home cage (n=5 sections from 4 mice), tone only (n=4 sections from 2 mice) and tone + shock (n=17 sections from 8 mice) conditions. Significantly more cholinergic neurons were reactivated following tone-shock pairings (Kruskal-Wallis p<0.01). Tone-shock compared to tone only (***, p=0.0017, Tukey corrected.) *See also Figure S2.

We performed a recall session (tone alone) 72 hours later and sacrificed the mice ∼2.5h following recall. We quantified basal forebrain cholinergic neurons for mCherry (neurons active during the training window) and GFP (cells activated during the recall session) expression after either training (Session 1;**Figure 2C**), or recall (Session 2; **Figure 2D**). Few cholinergic neurons were activated in the home cage or in response to tone alone (**Figure 2C**, home cage and tone). Significantly higher numbers of cholinergic neurons expressed mCherry following the tone-shock pairings (**Figure 2C**; Main effect p<0.01; tone-shock vs. tone only, p=0.004). These results parallel assays of ACh release in the BLA (**Figure 1**).

We next quantified the number of mCherry+/GFP+ (double positive) neurons following Session 2 (e.g. yellow arrow, **Figure 2B**). Significantly greater numbers of double positive cholinergic neurons were seen following the complete (tone + shock followed by tone recall) associative threat learning paradigm compared to mice that underwent training without shocks (**Figure 2D**, p=0.0017).

### Reactivation of cholinergic neurons activated by training is required for learned behavioral responses

BLA-projecting cholinergic neurons acquire tone responsiveness following associative threat learning (**Figure 1**) and a discrete population of NBM cholinergic neurons activated during tone-shock pairing, are reactivated during the recall session (**Figure 2**), important properties of memory engram neurons. If these cholinergic neurons are indeed part of a threat memory engram, then their reactivation should be required for generation of learned threat responses. To specifically prevent reactivation of cholinergic neurons in response to tone, we expressed the inhibitory, designer receptor hM4Di, in an activity dependent, Cre dependent manner in NBM/SI_p_ cholinergic neurons (ADCD-hM4Di; **Figure S2**) and subjected these animals to the threat learning paradigm (**Figure 3**). Animals were off Dox-chow during the training session, immediately placed back on Dox-chow and then tested for tone recall after 72 hr. ADCD-hM4Di and sham operated control animals were injected with clozapine (0.1 mg/kg) during the retrieval session to selectively silence the population of NBM/SI_p_ cholinergic neurons that were previously activated during training (**Figure 3A**). The control animals displayed normal freezing behavior in response to tone (**Figure 3B**; pre-tone vs. tone p=0.049, pre-tone vs. post-tone p=0.008). ADCD-hM4Di animals, in contrast, showed no freezing in response to the tone (**Figure 3B**; pre-tone vs. tone/post-tone, p>0.2), and overall showed decreased freezing behavior compared to controls (controls – black, hM4Di – red: # p=0.016), indicating that the reactivation during recall of training-activated cholinergic neurons in the NBM/SI_p_ was required for the expression of learned threat response behavior.

**Figure 3.**
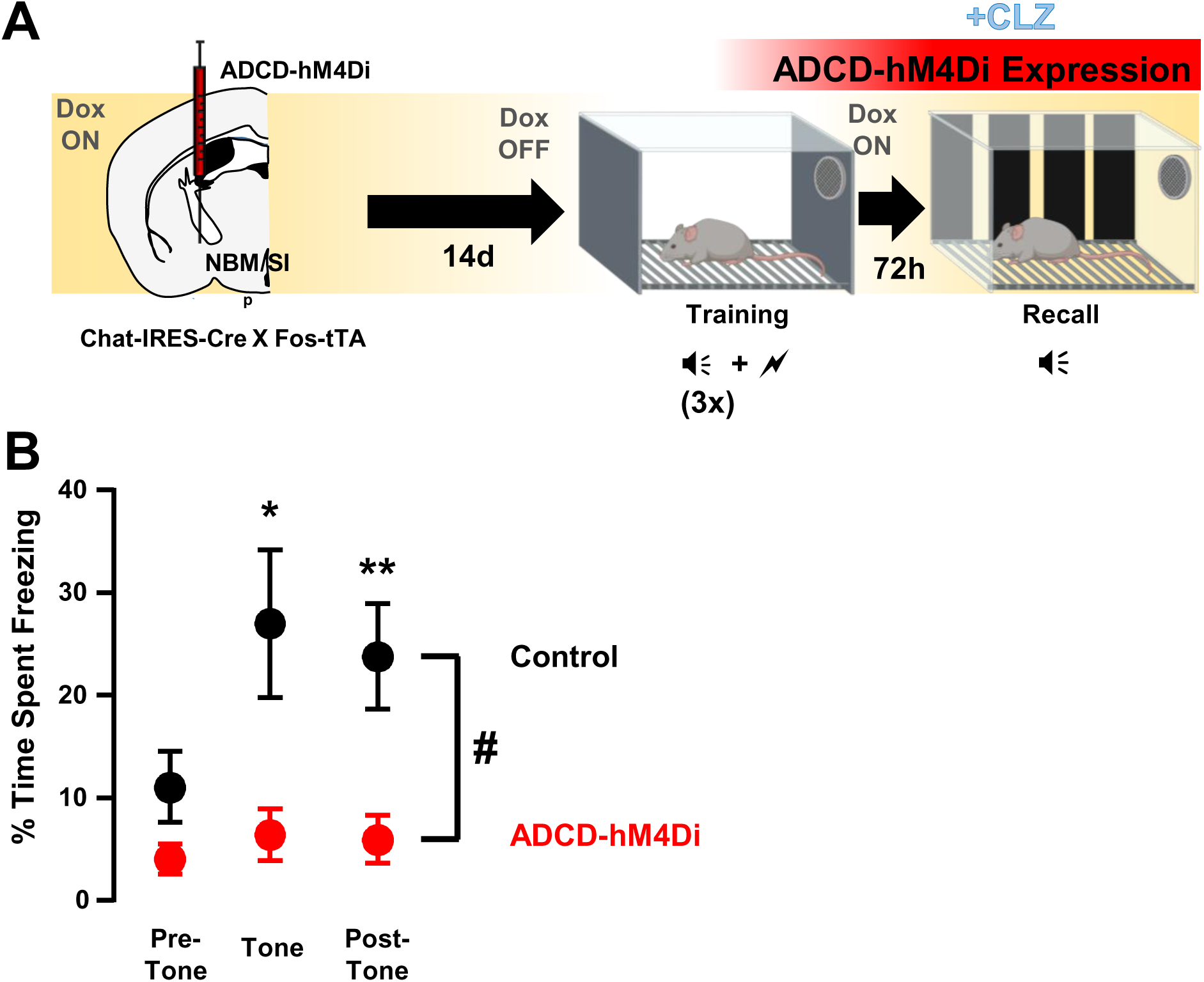
Re-activating NBM/SI_p_ cholinergic neurons is necessary for threat memory retrieval. **A.** ADCD-hM4Di was injected into the NBM/SI_p_ of Chat-IRES-Cre x Fos-tTA mice. Animals underwent training on regular chow (Dox off) to allow hM4Di.mCherry expression in training-activated cholinergic neurons; recall was tested on dox chow (Dox on) 72h following training. Clozapine (CLZ) was injected 10min before the recall session to prevent reactivation of cholinergic neurons that had been activated during training. **B.** Freezing behavior during recall following inhibition of training-activated cholinergic neurons in the NBM/SI_p_. Control (**black**, n= 8 mice) and hM4Di (**red**, n= 7 mice) groups. There were significant differences between pre-tone v. post-tone /tone freezing for controls (pre-tone v. tone *, p=0.0497; pre-tone v. post-tone **, p=0.0078; Tukey) and between control and hM4Di groups across the recall session (RM two-way ANOVA (GLM) TimexCondition #, p=0.0356).

### Recall-induced activation of NBM/SIp cholinergic neurons corresponds with the degree of threat response behavior

Both the level of freezing during the recall session, and the number of candidate threat engram cholinergic neurons varied considerably between animals. To address whether there was a relationship between the extent of freezing and the engagement of the cholinergic neurons, we labeled cholinergic neurons activated during the recall session with ADCD-mCherry (on dox during training, dox off during recall; **Figure 4A**). We divided animals into two groups based on behavior: low responders were defined as animals that showed no significant change in freezing between the pre-tone and the tone periods during recall (i.e. they did not display tone induced freezing); high responders showed an average 3-fold increase in freezing to tone compared to the pre-tone period (**Figure 4B**, p=0.005). We then quantified the fold change in the numbers of mCherry+ neurons in each group relative to home cage (**Figure 4C**). There was no difference in mCherry expression in low responders compared to the homecage group (fold change ∼ 1), but there was a 3-fold increase in the high responders (high responders vs. low responders, p=0.012, **Figure 4C**).

**Figure 4.**
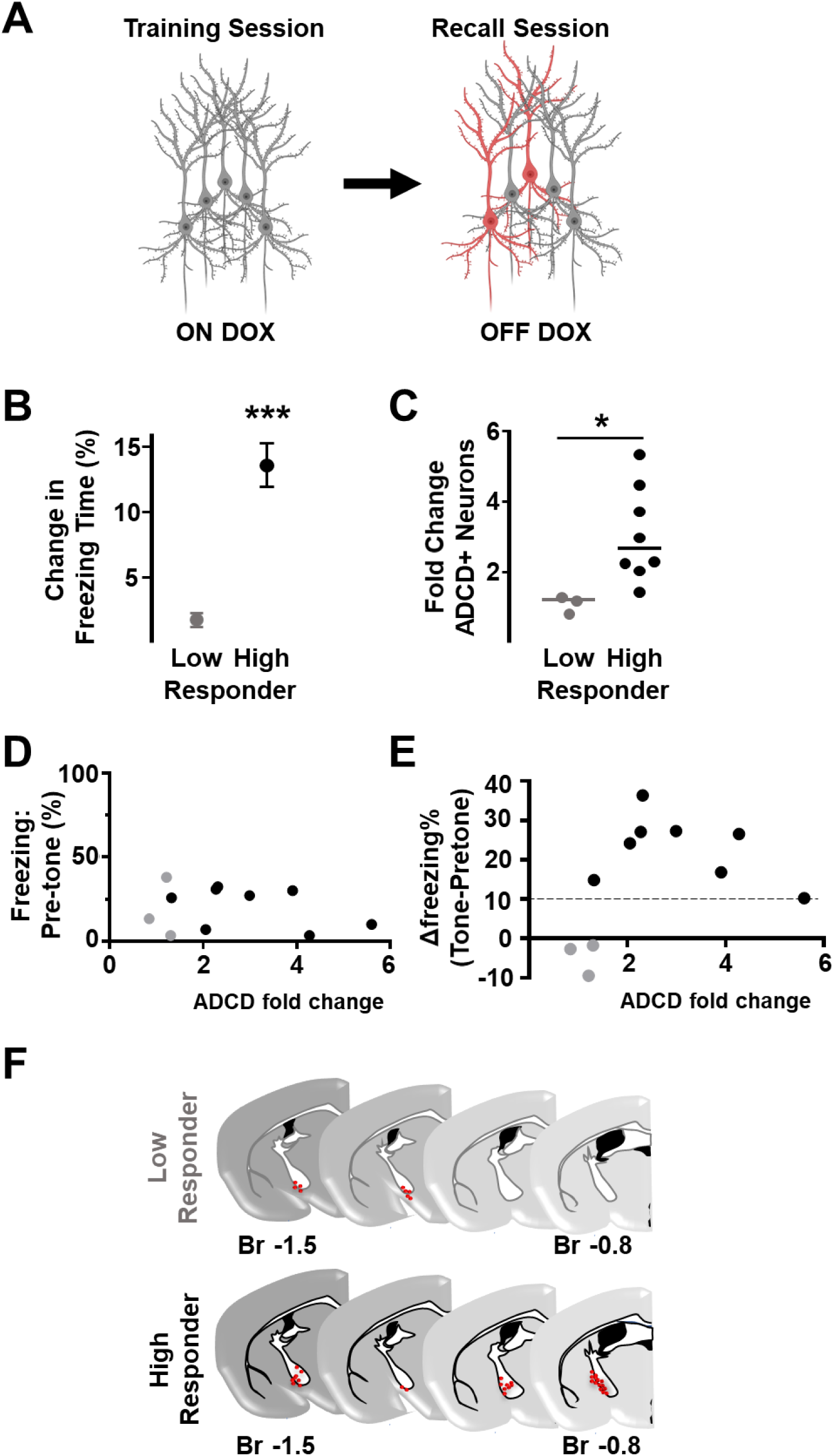
Activation of cholinergic neurons in the anterior NBM/SI_p_ is proportional to the conditioned response during threat-memory recall (see also Figure S3). **A.** Animals injected in the NBM/SI_p_ underwent training on Dox and recall off Dox to label recall activated NBM/SI_p_ cholinergic neurons. **B.** Quantification of freezing responses in low and high responders (See Methods for stratification criteria. n=5 low-responder, n=8 high-responder) (change in percent time spent freezing during recall between the 30s pre-tone bin and during the 30s tone bin). High responders froze significantly more during the tone than during the pre-tone period (Mann-Whitney test ***, p=0.005). **C.** Quantification of change in number of cholinergic neurons activated (ADCD-mCherry+) in low or high responders relative to the home cage. The number of ADCD+ neurons differed significantly between low and high responders (Mann-Whitney test *, p=0.0121) (n=3 low responder, n=8 high responder). **D.** Relationship between pre-tone freezing and increase in ADCD+ cells during recall. Each dot represents a single animal. Gray – low responders (n=3), black – high responders (n=5). Pearson correlation R^2^ =0.045, p=0.52. **E.** Relationship between the change in freezing (Δ freezing = tone freezing – pre-tone freezing) and increase in ADCD+ cells in response to recall tone. Gray – low responders, black – high responders. Dotted line indicates 10% points change in freezing (threshold for high response; see methods). Nonparametric Spearman correlation r=0.618 p=0.0478 (fold change in freezing v. fold change in ADCD+ cells). **F.** ADCD+ cholinergic neurons were plotted along the A-P axis (coronal images adapted from Paxinos atlas). Clusters (red dots) of activated cholinergic neurons were present in the anterior NBM of high responders (bottom) but not in low responders (top). See also Figure S3.

Freezing during the pre-tone period did not correlate with numbers of ADCD-mCherry expressing cells (**Figure 4D**, Pearson R^2^ =0.045, p=0.52). Comparing the change from pre-tone to tone-induced freezing behavior with number of ADCD-mCherry expressing cells (**Figure 4E**) revealed a positive correlation between fold change in freezing and fold change in number of ADCD labeled cells (Spearman r=0.618, p=0.0478) and the existence of 2 populations of mice delineated by an increase in activation of NBM/SIp BFCNs: one in which increased freezing (>10%) was associated with increased ADCD labeling (>2-fold) and a second in which there was no association between changes in freezing and ADCD labeling.

Cholinergic neurons activated during recall in high responders were in anatomically distinct regions from those in either low responders or home cage (**Figure 4F**). In the low responders, the small number of ADCD-mCherry labeled neurons were found in caudal regions of the NBM/SI_p_ (∼Bregma −1.5; **Figure 4F-top**), whereas in high responders there was an additional population in more rostral aspects of the NBM/SI_p_ (∼Bregma −0.7/-0.8; **Figure 4F-bottom**). Thus, there was a discrete population of activated cholinergic neurons in the rostral NBM/SI_p_ that correlated with the response to learned threat. Remarkably, retrograde mapping of BLA-projecting cholinergic neurons using CAV_2_-DIO-hM4Di.mCherry delivered in the BLA of Chat-IRES-Cre mice, shows a similar distribution along the rostro-caudal axis of the NBM/SI_p_ (**Figure S3A**). Taken together these data support the presence of a threat memory engram in the BLA-projecting BFCN population in the NBM/SI_p_.

### Silencing BLA-projecting basal forebrain cholinergic neurons prevents activation of BLA neurons

BLA neurons activated by threat learning transiently express Fos, which is then re-expressed following threat memory retrieval in the same neurons. This re-expression has been used to identify engram cells in the BLA (Josselyn et al., 2015; Reijmers et al., 2007; Tonegawa et al., 2015).

To determine whether silencing BLA-projecting cholinergic neurons during training or recall affected the activation of BLA neurons, we injected the BLA of Chat-IRES-Cre mice with CAV_2_-DIO-hM4Di.mCherry and AAV_9_-CaMKIIa-GCaMP6f (cav.hM4Di^BLA^ animals) or AAV_9_-CaMKIIa-GCaMP6f alone (control animals) (**Figure 5A**) (GCaMP6f was used as in injection site marker). mCherry expression was found predominantly in the NBM/SI_p_ identifying these cholinergic neurons as the major BLA-projecting population, followed by the VP/SI_a_ and a small contribution from the horizontal limb of the diagonal band of Broca (hDB) (**Figure 5A right**).

**Figure 5.**
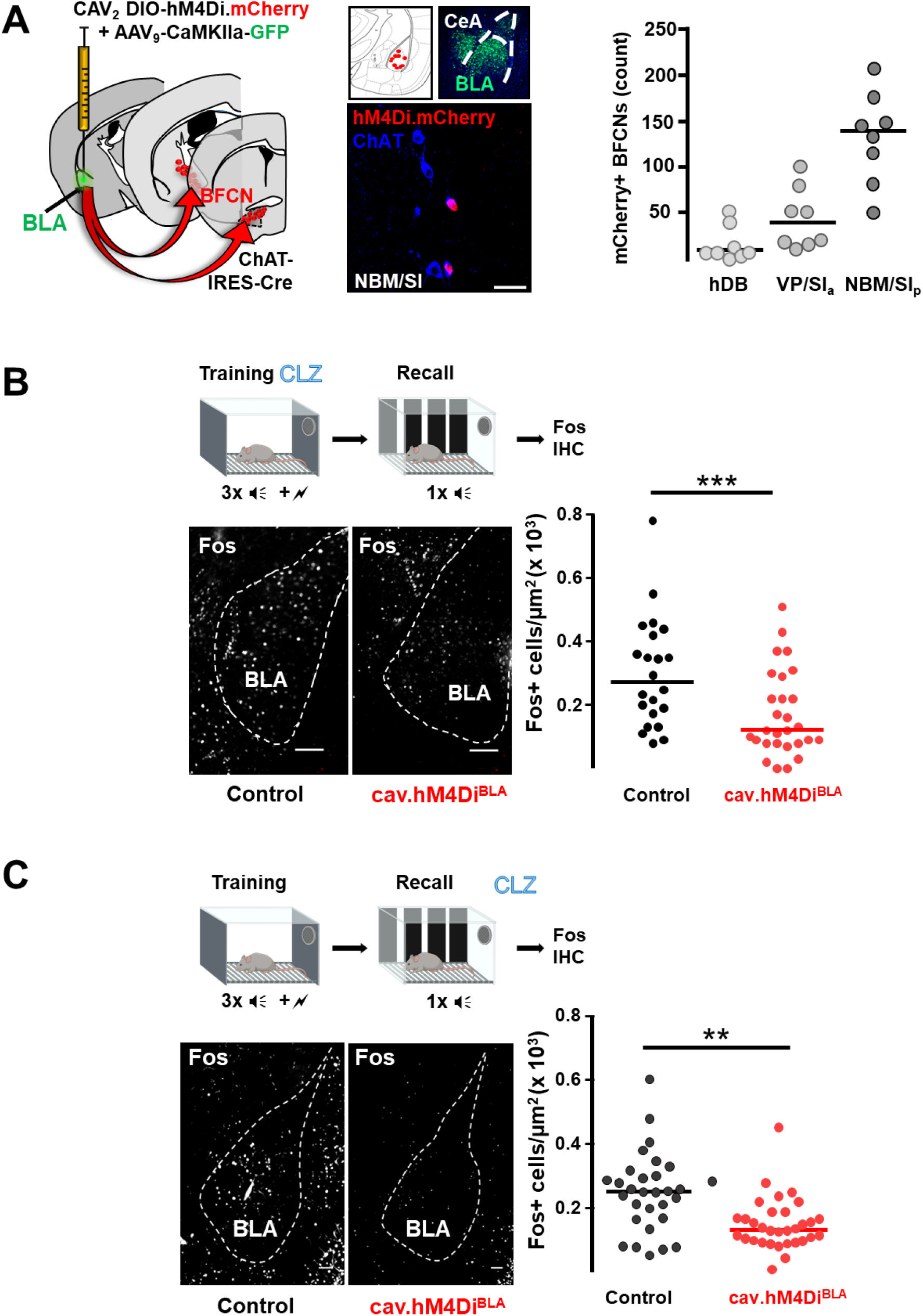
BLA-projecting cholinergic neuronal activity is required during training and recall for learned threat processing. (see also Figure S4) **A. Left**. Strategy for retrograde targeting of hM4Di DREADD to BLA-projecting cholinergic neurons. **Middle.** Re-localization of BLA injection sites (using P_CAMKIIα_-GCaMP6f to mark the injection site), and identification of retrogradely-labeled cholinergic neurons within the NBM/SI_p_ (right, image taken at A/P ∼ −1.34 from Bregma scale bar = 50µm). **Right.** Quantification of hM4Di-expressing cholinergic populations (mCherry+) across the basal forebrain (n=8 mice, 56-80 sections/animal). **B.** DREADD inhibition of BLA-projecting cholinergic neurons during **TRAINING** reduced BLA Fos immunoreactivity following recall. **Right.** Fos+ cell density in BLA control (black) vs. CAV_2_-DIO-hM4Di.mcherry (red) (n= 7 mice/group, 22 sections control vs. 28 sections hM4Di. Each dot corresponds to one section). Mann-Whitney test ***: p=0.0043. Lines represent median for each group. **C.** Silencing BLA-projecting cholinergic neurons during **RECALL** reduced BLA Fos immunoreactivity following recall. BLA images following Fos immunostaining. Fos+ cell density in BLA between control (black) vs. CAV_2_-DIO-hM4Di.mcherry (red) injected animals (n=3 mice/group, 30 sections control vs. 32 sections hM4Di. Each dot corresponds to one section. Mann-Whitney test **: p=0.005). See also Figure S4.

We injected cav.hM4Di^BLA^ or control mice with clozapine (CLZ) 10 min prior to initiating cue-conditioned threat learning (**Figure 5B**) **<u>or</u>** 10 min prior to the memory recall session (**Figure 5C**). In both cases mice were sacrificed 45-60 min following recall and assessed for Fos immunoreactivity (IR) in the BLA. Mice in both control groups showed equivalent density of Fos-IR cells indicating that 0.1mg/kg clozapine did not alter Fos expression (**Figure 5B & 5C; Control groups-black circles**). Silencing BLA-projecting cholinergic neurons **<u>during training alone</u>** blunted recall-induced activation of BLA neurons (**Figure 5B**: control v. cav.hM4Di^BLA^ p=0.0043). Silencing BLA-projecting cholinergic neurons **<u>during recall alone</u>** also reduced recall-induced activation of BLA neurons (**Figure 5C**: control v. cav.hM4Di^BLA^ p=0.005). Thus, activity of BLA-projecting cholinergic neurons was required during **<u>both</u>** learning and recall to successfully induce Fos expression in BLA neurons; preventing either significantly reduced the number of Fos+ BLA neurons.

Differences in recall-induced Fos expression between control and cav.hM4Di^BLA^ mice were maximal in rostral portions of the BLA (between bregma −0.8mm to −1.4mm) (**Figure S4B**). This region of the rostral BLA has been shown to contain genetically distinguishable neurons that are preferentially activated by aversive stimuli and preferentially project to the CeC, a region known to drive freezing behavior (Kim et al., 2016; Kim et al., 2017). We examined the CeC of mice in which BLA-projecting BFCNs were silenced during recall and found that there was a significant difference in Fos+ cell density between control and cav.hM4Di^BLA^ mice (**Figure S4C** control v. cav.hM4Di^BLA^ p=0.0012). Injection of CAV_2_-DIO-hM4Di into the BLA showed some spread of virus to the CeA (**Figure 5A middle**). To investigate whether silencing of any CeA-projecting cholinergic neurons could have mediated the effects on CeC Fos-IR, we injected CAV_2_-DIO-hM4Di directly into the CeA and silenced CeA-projecting cholinergic neurons during recall by injecting mice with clozapine 10 min prior to recall. There were no significant differences in freezing behavior between control and cav.hM4Di^CeA^ mice (**Figure S4H**). Thus, specifically silencing cholinergic input to the BLA altered activation of BLA circuits involved in execution of defensive behaviors. Given these findings, we examined freezing behavior in these groups of mice (control vs. cav.hM4Di^BLA^).

Behaviorally, both groups of control mice (clozapine during training or clozapine during recall) displayed significant freezing in response to the recall tone (**Figure S4D and S4E**; **black circles**, **S4D** pre-tone to tone p=0.03; **S4E**: p=0.001); whereas cav.hM4Di^BLA^ mice that received clozapine during training did not freeze to the tone and cav.hM4Di^BLA^ mice that received clozapine during recall had a significantly blunted freezing response to the tone (**Figure S4D & 4E**; **red circles**, CAV.hM4Di^BLA^ vs. control, clozapine during training (**Figure S4D**) pre-tone to tone p=0.22; clozapine during memory recall (**Figure S4E**), p=0.06). Since hM4Di-mediated silencing can last several hours (Ryan et al., 2015), we inhibited BLA-projecting cholinergic neurons immediately after the training session (i.e. during early consolidation). This did not alter the freezing response during the recall session (**Figure S4F**). Thus, the activity of BLA projecting cholinergic neurons during tone-shock pairing and during recall, is essential for associative learning.

Mice in which cholinergic input to the BLA was silenced during recall still showed a modest increase in freezing response to the tone as a group (albeit significantly less than control mice) (**Figure S4E red circles**, pre-tone to tone p=0.06). To investigate whether this increase was driven by a subset of mice, we analyzed the proportion of high and low responding mice in each group of mice. We found that under control conditions (i.e. no cholinergic silencing), 80-90% of the mice were high responders and that silencing BLA-projecting cholinergic neurons **<u>during training</u>** resulted in 100% of the mice being low responders (**Figure S4G** control v. cav.hM4Di^BLA^ inhibition during training). On the other hand, silencing BLA-projecting cholinergic neurons **<u>during recall</u>** resulted in ∼50% of the mice being low responders (**Figure S4G** control v. cav.hM4Di^BLA^ inhibition during recall). Thus, silencing BLA-projecting cholinergic neurons only during recall resulted in an all-or-none behavioral phenotype.

### Cholinergic neurons activated during threat memory recall have altered intrinsic excitability

Changes in excitability of neurons have been consistently associated with the threat memory engram (Cai et al., 2016; Pignatelli et al., 2019; Rashid et al., 2016; Zhang and Linden, 2003; Zhou et al., 2009). We asked whether cholinergic neurons activated during memory recall differed in their intrinsic excitability compared to non-activated cholinergic neurons. Two to three hours after the recall session, we prepared acute brain slices from Fos-tTA/GFP mice for electrophysiological recording of activated (Fos-GFP+) and non-activated (Fos-GFP-) NBM/SI_p_ cholinergic neurons (**Figure 6A**; cholinergic neuron identity was determined by post-recording single cell RT-PCR). Cholinergic neurons that were Fos+ following the recall session differed significantly from Fos-cholinergic neurons and from cholinergic neurons from home cage mice (**Figure 6B&C**). Properties that showed significant differences included: action potential (AP) half-width, rheobase and maximum firing rate (**Figure 6D**; half-width: HC vs. Fos+ p=0.0006, Fos- vs. Fos+ p=0.021; **Figure 6E**; rheobase: Fos- vs. Fos+ p=0.023; **Figure 6F**; max firing rate: HC vs. Fos+ p=0.005, Fos- vs. Fos+ p=0.0034) as well as latency to fire (**Figure S5C**; latency: HC vs. Fos+ p=0.0004) and afterhyperpolarization (AHP) amplitude (**Figure S5E**, HC vs. Fos+ p=0.0041). AP amplitude and resting membrane potential did not differ.

**Figure 6.**
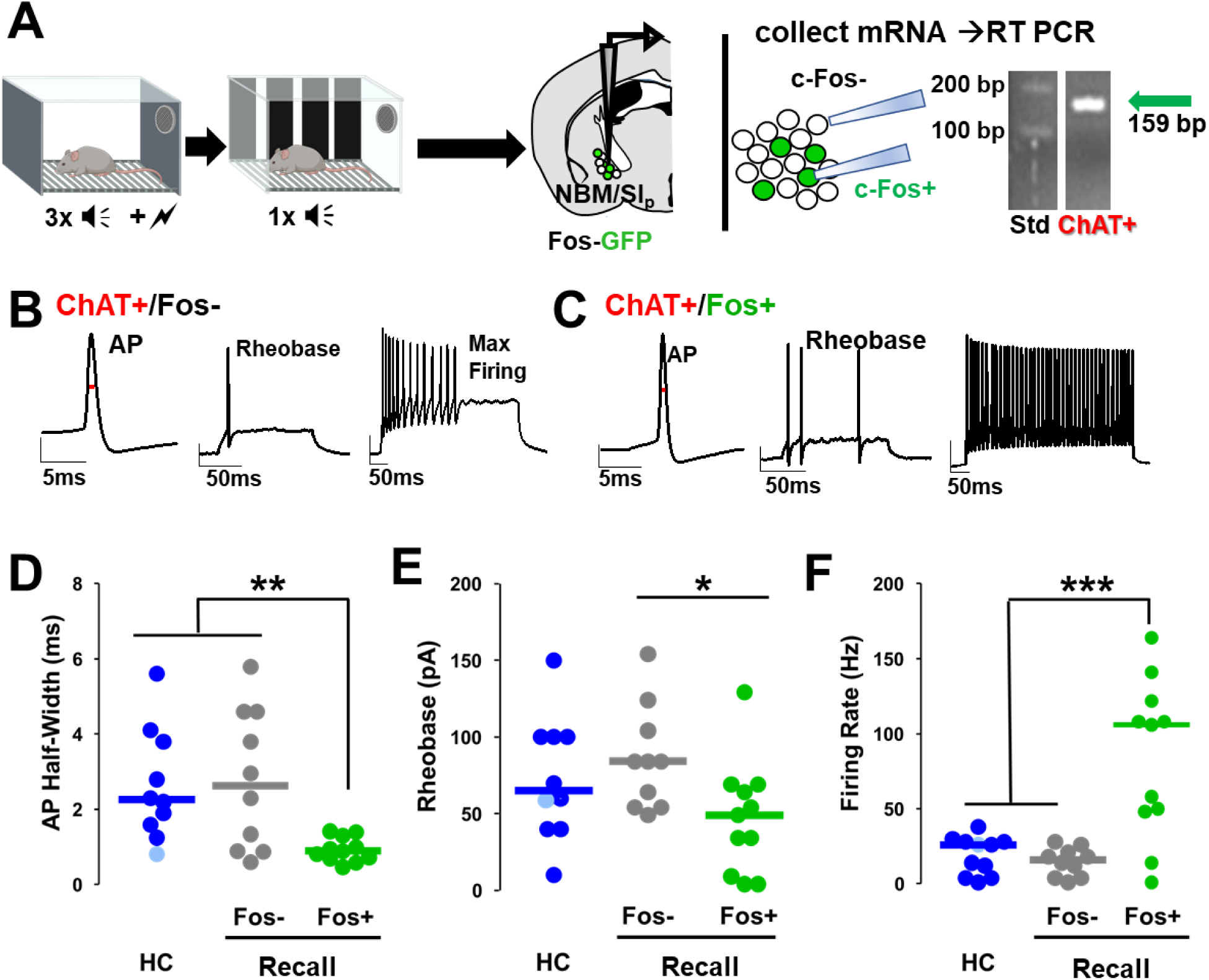
NBM/SI_p_ cholinergic neurons show increased intrinsic excitability following threat memory recall (see also Figure S5) **A.** Schematic of electrophysiological profiling of activated (Fos+) vs non-activated (Fos-) neurons with *post-hoc* identification of cholinergic identity by single cell RT-PCR. **B.** Representative traces following injection of current into a Fos-NBM cholinergic neuron (**ChAT+/Fos-**). Red line denotes AP-half width measurement. **C.** Representative traces following current injection into a Fos+ NBM cholinergic neuron (**ChAT+/Fos+)**. Red line denotes AP-half width measurement. **D, E, F:** Population data (dot plot + line at median) for the electrophysiological properties of *post-hoc* identified cholinergic neurons. Analyses assess active membrane properties including action potential (AP) half-width (D), rheobase (E) and the maximal firing rate in response to prolonged steady state depolarization (F), from home cage (HC; n= 11 ChAT+ neurons from 11 mice), and following recall to tone alone (Fos-: n= 10 ChAT+ neurons from 5 mice; Fos+: n= 11 ChAT+ neurons from 6 mice). **D:** Kruskal-Wallis tests; AP half-width: **:p=0.0054 (Dunn’s Corrected p-values: HC vs. Fos-: p = 0.8971, HC vs. Fos+: p = 0.0006, Fos- vs. Fos+: p = 0.0206) **E**: Rheobase: *: KW=p=0.05 (Dunn’s Corrected p-values: HC vs. Fos-: p = 0.0792, HC vs. Fos+: p = 0.3897, Fos- vs. Fos+: p = 0.0228) **F**: Max firing rate: ***: p=0.0019. (Dunn’s Corrected p-values: HC vs. Fos-: p = 0.6959, HC vs. Fos+: p = 0.0005, Fos- vs. Fos+: p = 0.0034) *See also Figure S5.

We also compared the firing rate of cholinergic neurons in home cage mice with those expressing Fos 2 hr after training or after recall (measured from 2 hr through 5 days after the recall session **Figure S5G&H**). We found no differences in firing rate between home cage cholinergic neurons and cholinergic neurons that expressed Fos after training. Further, the increase in maximal firing rate seen after recall returned to baseline within 3-5 d (compared to Recall D0, p<0.01 for all). Together, these results indicate cholinergic neurons activated during threat memory recall underwent biophysical changes that increased their excitability, a finding that, combined with their activation-reactivation profile, their requirement for the activation of the BLA engram, and their requirement for display of learned threat behaviors, argues that this population of cholinergic neurons is a critical component of the learned threat memory engram.

### Distinct subsets of BLA-projecting cholinergic neurons differentially contribute to learned vs. innate threat processing

Having established a role for basal forebrain cholinergic neurons in a learned threat paradigm, we asked whether these neurons also contribute to innate threat responses. We used predator odor exposure as a model for innate threat processing (Blanchard and Blanchard, 1990). To examine whether innate threat responses engage basal forebrain cholinergic neurons and/or require the activity of basal forebrain cholinergic neurons, we exposed Fos-tTA/GFP mice to a mountain lion (Mt. Lion) urine wetted gauze pad (**Figure 7A**). Exposed mice showed increases in active and passive defensive behaviors compared to mice exposed to a saline wetted pad, including freezing (**Figure 7A**, p=0.028), avoidance (**Figure S6A**, p=0.002) and defensive digging (**Figure S6A**, p=0.026).

**Figure 7:**
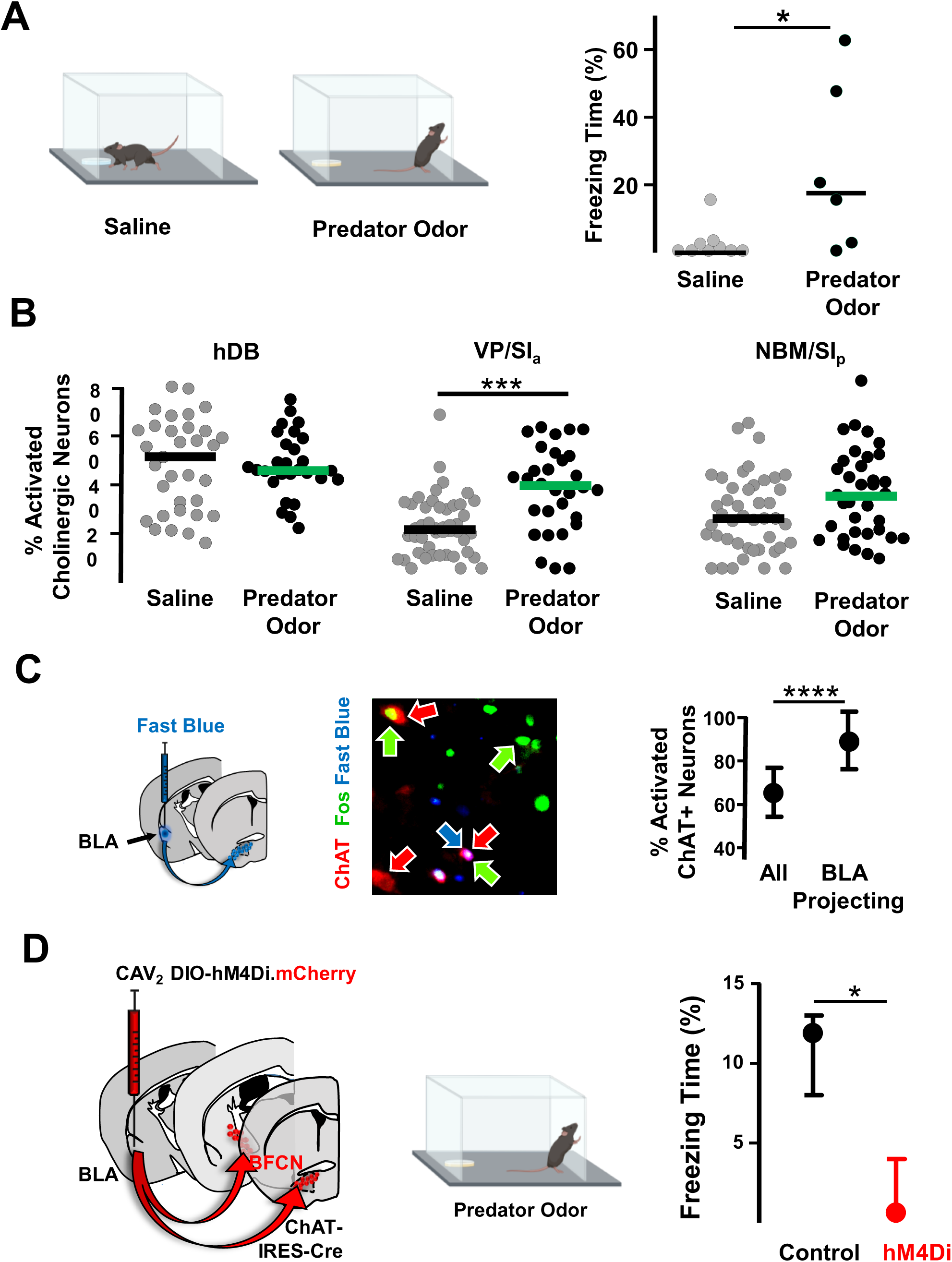
Distinct subsets of BLA-projecting cholinergic neurons differentially contribute to learned vs. innate threat processing (see also Figure S6) **A.** Mice were placed in chambers containing a gauze pad spotted with either saline or with mountain lion urine (predator odor). Defensive behaviors were monitored for 5 min. Animals froze significantly more in the presence of predator odor than saline (Mann-Whitney *, p= 0.028). **B.** Quantification of activated cholinergic neurons (Fos-GFP+,ChAT+) in basal forebrain nuclei that project to the BLA in response to predator odor grouped by region. Predator odor exposure increased the number of activated cholinergic neurons in the VP/SI_a_ (Mann-Whitney****: p <0.0001), but not in the hDB (p = 0.9) or NBM/SI_p_ (p=0.56).). Data points are from individual sections from n=7 control and n=6 odor exposed animals. **C.** Fast blue was injected into the BLA to retrogradely label BLA projecting neurons prior to odor exposure. After exposure to predator odor sections from the basal forebrain were immunostained with antibodies recognizing ChAT and Fos and the numbers of activated cholinergic neurons were counted (ChAT+Fos+/total ChAT+). In the VP/SI_a_ overall ∼65% of cholinergic neurons were activated, and over 90% of BLA-projecting cholinergic neurons were activated (ChAT+ red, Fos+ green, Fast Blue blue, n=3 mice). Data are displayed as mean % of activated cholinergic neurons per section +/- SD (n=15 sections from 3 mice). **** indicates preferential activation of BLA-projecting VP/SI_a_ cholinergic neurons (p=0.0001). **D.** Chat-Cre mice injected in the BLA with a control virus (AAV-CAMKIIa-GFP) alone (control) or in combination with CAV_2_-DIO-hM4Di were exposed to predator odor following injection with clozapine (CLZ). Freezing behavior was measured during a 5 min exposure (mean ±SD; control – black, hM4Di – red). Silencing BLA-projecting cholinergic neurons significantly blunted the freezing response (Mann-Whitney *: p=0.019; control: n=6; hM4Di: n=4 mice). See also Figure S6.

Next, we quantified the number of cholinergic neurons expressing Fos-GFP after saline or predator odor exposure (**Figure 7B**; Fos-GFP+ChAT+). The number of Fos-GFP expressing cholinergic neurons was significantly elevated in the predator odor exposed group in the VP/SI_a_ (**Figure 7B**, p<0.0001), but not the hDB (**Figure 7B**, p=0.9) or NBM/SI_p_ (**Figure 7B**, p= 0.25).

VP/SI_a_ cholinergic neurons formed the second largest source of cholinergic input to the BLA in our retrograde mapping experiments (**Figure 5A**), and cholinergic lesions of the BLA have been shown to reduce freezing behavior in rats exposed to cat hair (Power and McGaugh, 2002). Since VP/SI_a_ cholinergic neurons were found to be activated during predator odor exposure, rather than NBM/SI_p_ or hDB cholinergic neurons, we asked if the BLA-projecting pool of VP/SI_a_ cholinergic neurons was activated by predator odor exposure. We injected the retrograde tracer Fast Blue into the BLA of Fos-GFP mice and then exposed them to either saline (control) or Mt. Lion urine (**Figure 7C left**). Fast Blue labeled approximately 30% of ChAT immunoreactive neurons located in the VP/SI_a_ (data not shown). Nearly the entire subset of BLA-projecting VP/SI_a_ cholinergic neurons were also Fos-GFP+ (**Figure 7C right**, All vs. BLA-projecting, p=0.0001).

To determine whether activity of these BLA-projecting cholinergic neurons was necessary for animals to show a defensive response to predator odor, we used CAV_2_-DIO-hM4Di to silence BLA-projecting cholinergic neurons. Silencing during predator odor exposure resulted in significantly less freezing compared to control mice (**Figure 7D right** control v. cav.hM4Di^BLA^ p=0.019) but did not alter avoidance or digging behavior (**Figure S6B**). Taken together, these data support the conclusion that activity of BLA-projecting VP/SI_a_ cholinergic neurons is critical for maintaining normal passive defensive behaviors in response to innate threat processing.

Finally, we asked whether VP/SI_a_ BFCNs also contribute to freezing behavior in response to learned threats. We delivered AAV_9_-DIO-hM4Di.mCherry or AAV_9_-DIO-eCFP into the VP/SI_a_ or NBM/SI_p_ of Chat-IRES-Cre mice (**Figures S6C & S6D**). Both hM4Di and eCFP animals were injected with clozapine prior to the recall session. Animals in which NBM/SI_p_ cholinergic neurons were silenced during the recall session did not freeze in response to tone (**Figure S6C**, control, black: pre-tone to tone p=0.0049; aav.hM4Di^NBM^, red: pre-tone to tone p=0.37), whereas animals in which VP/SI_a_ cholinergic neurons were silenced froze to the same extent to the tone as the clozapine injected eCFP animals (**Figure S6D**, p=1.00).

In sum, two distinct paradigms of threat exposure (learned and innate) require activity in BLA-projecting basal forebrain cholinergic system, although the populations engaged are distinct

## Discussion

A small number of sparsely distributed neurons across the basal forebrain provide extensive cholinergic innervation to most of the brain. Though few in number, these cholinergic neurons and their widespread projections play a critical role in modulating cognitive processes (Ballinger et al., 2016; Záborszky et al., 2018). Transient optogenetic stimulation of ACh release is necessary and sufficient to induce long term potentiation of glutamatergic transmission at cortico-BLA synapses (Jiang et al., 2016). What remains unclear is whether the cholinergic system encodes stimulus-specific information, or whether it is generally recruited to modulate salience across long timescales. To begin addressing these questions, we monitored ACh release in the BLA during threat learning and retrieval, anatomically mapped and electrophysiologically characterized behaviorally relevant BFCNs, and investigated the contribution of different subsets of BFCNs to threat response behaviors. Taken together, our results support the presence of a cholinergic component of the threat memory engram.

### Basal-Forebrain Cholinergic Neurons “Learn” to Respond to the Conditioned Stimulus

In this study we used a genetically encoded ACh sensor (GRAB_ACH3.0_) to monitor endogenous ACh release in the BLA during threat learning and recall. First, we found that foot-shock rapidly and reliably evoked ACh release, in line with previous observations (Jing et al., 2020). It has previously been proposed that BFCN responses diminish to expected outcomes during reward learning (Hangya et al., 2015). In our 3X tone-shock pairing (fear learning) paradigm, where mice are naïve only at the presentation of the first tone (CS) and first shock (US), we found that responses to the first foot-shock were the fastest and had the highest amplitude when compared to responses to the second and third shocks (Figure 1B). When we examined responses to the tone (CS), we found that BFCNs have little to no response to a naïve, unexpected tone. Just three minutes after the first tone-shock pairing, we observed an increase in ACh release in response to the second tone, without change to the kinetics of the release. This response was further amplified and presented with shorter latency upon third tone presentation. These time scales match previously observed time scales for activity-dependent plasticity of glutamatergic transmission *in vivo,* and support plasticity of auditory-evoked potentials in basal-forebrain cholinergic neurons upon CS-US pairing (Guo et al., 2019; Stephenson-Jones et al., 2020). Importantly, we did not observe significant tone induced changes to ACh release in the BLA when tone was not paired with a shock (**Figure S1**). When mice were exposed to the conditioned tone in a novel environment 24h later, we observed robust ACh release in the BLA compared with the naïve tone (**Figure 1D**). This enhancement of ACh release supports the notion that BFCNs undergo physiological changes which allow robust responsiveness to previously naïve inputs.

### Cholinergic Modulation of Associative Threat Learning

Recent work examining mechanisms of learning and memory in the BLA have revived Semon’s theory on memory engrams: learning must result in lasting biophysical changes that form the substrate for retrieval of the learned experience (Semon, 1921; Tonegawa et al., 2015). In the BLA, several molecular changes occur in response to learning of CS-US associations, including new gene expression and protein synthesis (Sears et al., 2014). Calcium induced signaling cascades initiated as a consequence of “suprathreshold” synaptic activity (Greer and Greenberg, 2008), induce expression of immediate early genes (IEG). IEG expression has been used as a proxy to identify “activated” neurons. To qualify as an “engram neuron,” a neuron must be activated not only during learning, but upon retrieval of that memory (Tonegawa et al., 2015). This has been previously observed in BLA neurons that show fos expression following CS-US pairings and following recall of those memories in response to CS alone.

Using DREADD-mediated silencing of BLA-projecting BFCNs we found that silencing BLA-projecting cholinergic neurons either during training or during recall resulted in loss of freezing behavior as well as significant reductions in density of Fos+ neurons in the BLA following recall (**Figure 5**). The reduction of Fos expression in the BLA following recall when BLA projecting cholinergic neurons were silenced during training suggests that cholinergic signaling in the BLA is necessary for establishing the BLA threat memory engram. Additionally, we found silencing cholinergic activity specifically during recall also disrupted recruitment of BLA neurons and subsequent expression of freezing behavior. Taken together, these data support the critical role of BLA-projecting cholinergic neurons in formation of the BLA engram.

We have previously demonstrated that activation of presynaptic acetylcholine receptors can induce sustained potentiation of glutamate release (Jiang et al., 2013; Jiang et al., 2016; McGehee et al., 1995; Zhong et al., 2017; Zhong et al., 2008; Zhong et al., 2013, 2015). Jiang et al. (Jiang et al., 2016) demonstrated that the increased presynaptic release mediated by ACh was dependent on nicotinic acetylcholine receptors (nAChRs) located on presynaptic glutamatergic terminals in the BLA, and that nAChR activation in the BLA was necessary for acquisition of conditioned threat memories (Jiang et al., 2016). Importantly, it has been shown that BLA neurons recruited during memory recall exhibit increased presynaptic glutamatergic activity (Nonaka et al., 2014). Based on these findings, we propose that silencing BLA-projecting cholinergic neurons during threat learning or during recall results in loss of Fos expression due to alterations in glutamatergic transmission, and subsequent disruptions to the formation and/or recruitment of the BLA engram.

### A Cholinergic Component in the Associative Threat Memory Engram

We next sought to investigate whether NBM/SI_p_ BFCNs, the primary cholinergic input to BLA, were activated during threat learning and reactivated during memory recall. We found that cholinergic neurons in the NBM/SI_p_ were not activated following naïve tone or repeated presentation of tone not paired with a shock but were activated in response to tone-shock pairings (Fig 2). We found that the *same* cholinergic neurons activated during training were activated upon presentation of the now conditioned tone during recall. Importantly, selectively blocking the cholinergic engram by preventing reactivation during recall of the NBM/SI_p_ BFCNs that had been activated during training, was sufficient to block the freezing response to conditioned tone recall (Fig 3). Further, activation of NBM/SI_p_ BFCNs during recall correlated with freezing response (Fig 4) only when the tone was previously paired with a shock.

We found an interesting relationship between anatomical distribution of activated cholinergic neurons in high- and low-responding mice in the rostral portions of the NBM/SI_p_, a region with a large proportion of BLA-projecting cholinergic neurons (Fig S3). It is important to mention that at the relevant bregma locations (Bregma −0.7/-0.8 mm), BLA-projecting BFCNs are intermingled with ventrolateral orbitofrontal (VO/LO) and medial prefrontal cortex-projecting BFCNs (Gielow and Zaborszky, 2017). We cannot rule out an additional role for these cholinergic neurons in threat processing.

### Changes in excitability of Fos+ cholinergic neurons

It has been proposed that alterations to synaptic weights and changes in ionic conductance resulting from learning-induced transcriptional programs allow for increased response fidelity during memory retrieval (Yap and Greenberg, 2018). This is supported by studies where blocking new protein synthesis following learning results in loss of long-term memory (Hernandez and Abel, 2008). Based on this fundamental understanding of interactions between neural plasticity and memory, it can be proposed that a memory trace should be detectable during retrieval in a neuron: either as a pattern of activity that supports expression of retrieval behavior, or as a change in excitability that might precede synaptic plasticity. To assess whether such changes were present in recruited cholinergic neurons following memory retrieval, we recorded properties of neuronal excitability from activated NBM/SI_p_ BFCNs (Fos+) and compared them with Fos-BFCNs recorded from the same brain slices. We found that activated (Fos+) NBM/SI_p_ cholinergic neurons showed increased excitability following threat memory recall, which lasted for several hours following recall but eventually returned to baseline. This finding is in line with previous reports of learning-associated changes in electrical properties, which are found shortly after recall, but disappear at later time points despite the persistence of the learned behavior (Moyer et al., 1996; Pignatelli et al., 2019). These changes were not present in Fos-GFP+ cholinergic neurons immediately following training which were indistinguishable from cholinergic neurons in the homecage control group or other Fos-GFP-cholinergic neurons. Thus, the changes in electrical properties we observed were specific to recall-activated cholinergic neurons. It is possible that Fos-induced transcription and de novo protein synthesis drive changes to intrinsic properties, and thus do not arise for several hours following training. Within recall-activated cholinergic neurons we find changes in AP half-width, rheobase, and max firing rate. Interestingly, we find an increase in AHP amplitude measured after a single AP at rheobase and a trend toward reductions in duration in Fos+ cholinergic neurons. Common features of activated neurons previously reported include increases in firing rate, reductions in adaptation, reductions in AHP amplitude, reductions in duration of post-burst afterhyperpolarization (medium AHP), and synaptic alterations (Whitaker and Hope, 2018).

### Differential contribution of distinct BLA-projecting BFCNs in learned vs. innate threat processing

Amygdala microcircuits play an important role in the regulation of active vs. passive avoidance behaviors (Rickenbacher et al., 2017; Terburg et al., 2018). Our finding that silencing cholinergic input to the BLA resulted in a selective loss of threat-motivated freezing behavior supports potential specificity of cholinergic modulation within BLA microcircuits. Specifically, we found that BLA-projecting cholinergic neurons were necessary for freezing in response to a learned threat-associated cue, and for freezing in response to the innately threatening predator odor. Predator odor selectively activated BLA projecting, VP/SI_a_ cholinergic neurons and direct silencing of NBM/SI_p_, but not VP/SI_a_ cholinergic neurons attenuated learned threat induced freezing. Based on these data, we propose that distinct populations of BLA-projecting BFCNs control freezing in response to fundamentally distinct threatening situations (learned vs. innate).

Several studies support the notion that distinct cholinergic circuits regulate unique features of threat processing. Previous studies using pharmacological manipulations have suggested differences in cholinergic receptor contribution in the BLA during innate vs. learned freezing behavior. While learned freezing behavior was affected by blocking nicotinic acetylcholine receptors, innate freezing could be manipulated by blocking muscarinic acetylcholine receptors (Jiang et al., 2016; Power and McGaugh, 2002). Further, anatomical mapping has shown that VP/SI_a_ and NBM/SI_p_ BFCN projections travel distinct paths to reach the BLA and have overlapping yet separate terminations within BLA (Allen Brain Connectivity Atlas: experiment 161460864 vs. 478490368). Recent monosynaptic input mapping has also revealed that NBM/SI_p_ and VP/SI_a_ cholinergic neurons may receive distinct inputs (Gielow and Zaborszky, 2017; Hu et al., 2016). Whether these features interact to produce distinct cholinergic synaptic connectivity within the BLA, and whether these differences underlie the differential contribution of different subsets of BLA-projecting BFCNs to learned vs. innate defensive behaviors remains to be examined.

### What recruits cholinergic neurons into the engram?

Studies from the Josselyn lab (Dr.Sheena Josselyn & collaborators) have demonstrated that neurons in the amygdala compete to become part of the associative threat memory engram (Han et al., 2007). It has been proposed that this competition is driven by stochastic changes in intrinsic excitability gating incoming synaptic inputs (Han et al., 2007). This hypothesis also supports the idea that the memory engram forms at the intersection between stochastic changes in intrinsic excitability and precise cue-driven synaptic inputs. Whether these stochastic changes in excitability are truly stochastic in nature, resulting in the recruitment of a critical mass as a gate to the engram, or occur in a privileged subset of neurons (Shim et al., 2018; Zhang and Linden, 2003) requires further investigation.

Evidence supports the idea of the recruitment of a critical mass. For example, alterations to behavioral performance by inducing amnesia resulted in alterations to engram cell number, and infection efficiency-dependent behavioral changes occurred in response to exogenous alterations to biophysical properties of neurons (Rumpel et al., 2005; Ryan et al., 2015). Here we report that the number of engram enrolled NBM/SI_p_ cholinergic neurons correlates with behavioral performance, though it is unknown whether this subset is randomly recruited or predetermined. Several studies support extensive heterogeneity of cholinergic neurons across the basal forebrain. For example, cholinergic neurons within the same subregions have distinct projection paths and unique inputs. Furthermore, differences in electrical properties of cholinergic neurons have been described which allow them unique responses to the same input (Laszlovszky et al., 2020). Taken together, it becomes critical for future studies to examine the composition of the cholinergic engram to determine whether recruitment is biased to a privileged population or determined by random fluctuations to excitability.

Recent studies report enhancement of engram-to-engram cell synaptic connectivity following associative learning (Choi et al., 2018). We and others have shown that cholinergic neurons are rapidly activated by aversive stimuli such as footshock (Hangya et al., 2015; Jing et al., 2020), however during recall, we present only a tone to represent the previous association between the tone and shock. Presentation of the tone can engage prefrontal cortical and hippocampal processing, or even direct inputs that are known to exist from auditory cortical areas to BLA-projecting BFCNs, to reactivate cholinergic neurons (Bergstrom, 2016; Gielow and Zaborszky, 2017; Gilmartin et al., 2014). We hypothesize that footshock-induced synaptic input to the NBM/SI_p_ cholinergic neurons may, then, potentiate tone-related synaptic inputs via mechanisms such as heterosynaptic facilitation.

## Conclusion

The data presented in this study demonstrate recruitment of a modulatory component in an associate threat memory engram. We show that NBM/SI_p_ cholinergic neurons are recruited during threat memory acquisition and a subset of the same cholinergic neurons are reactivated during memory retrieval. We demonstrate that memory retrieval correlates with biophysical changes in NBM/SI_p_ cholinergic neurons including: emergence of a transcriptional response to the tone upon conditioning, alterations to electrical properties, and changes in ACh release in the BLA. We find that activation of NBM/SIp BFCNs during memory retrieval is directly correlated to the freezing response to the conditioned tone.

In addition to the BLA, cholinergic neurons in the NBM/SI_p_ region project to various limbic and sensory regions such as the lateral orbital cortex, cingulate cortex, somatosensory cortex, and mediodorsal thalamus. This raises the interesting possibility that the cholinergic engram can modulate various nodes of the threat memory engram circuit in conjunction with the amygdala allowing for coordinated retrieval of engrams across distributed networks. Such coordinated activation of distributed engrams has been recently demonstrated and may more closely recapitulate natural memory retrieval (Roy et al., 2019). Furthermore, functionally related regions have been shown to receive their cholinergic input from the same cholinergic nucleus (Zaborszky et al., 2015). We propose that engram-enrolled cholinergic neurons bind distributed engrams to produce stimulus-convergent, efficient memory retrieval. Future studies will test this hypothesis, investigating circuit and molecular mechanisms underlying engram-enrollment of basal forebrain cholinergic neurons.

## Acknowledgements

Part of this work conducted at Stony Brook University, NY was supported by NS022061 to LWR and DAT and MH109104 to LWR, DAT and MRP. Work conducted at the NIH was supported by the Intramural Research Programs of NINDS and NIMH. This work was also supported by NIDA grant DA14241, DA037566 and NIMH grant MH077681 to MRP. RBC was additionally supported by a NINDS Training Grant (T32) NS007224. We would like to thank Dr. Shaoyu Ge (Stony Brook University, NY) for providing reagents and insightful discussions aiding in the conceptualization of the project. P.R. would also like to thank Dr. Joshua Dubnau and Dr. Qiaojie Xiong (Stony Brook University, NY) for providing feedback and discussions on experiments presented in this manuscript. In additional we would like to thank Wendy Akmentin for expert technical assistance in data curation.

## Author Contribution

Conceptualization, D.A.T., L.W.R., P.R., M.R.P. Methodology, D.A.T., P.R., M.A., L.W.R., G.L.H., R.B.C., Y.L. Software, M.A., R.B.C. Validation, R.B.C. Formal Analysis, P.R., M.A., R.B.C. Investigation, D.A.T, P.R., M.A., R.B.C., L.J., G.L.H., C.A., S.W., A.J., C.Z., N.S.D. Resources, Y.L. Data Curation, M.A., R.B.C., P.R. Writing-Original Draft, P.R., M.A. Writing-Review & Editing, P.R., M.A., D.A.T., L.W.R., M.R.P., R.B.C. Visualization, M.A., P.R. Supervision, D.A.T., L.W.R., M.R.P. Project Administration, D.A.T., L.W.R., M.R.P. Funding Acquisition, D.A.T, L.W.R., M.R.P.

## Declaration of Interests

The authors declare no competing interests.

## STAR METHODS

### RESOURCE AVAILIBILITY

#### Lead Contact

Further information and requests for resources and reagents should be directed to and will be fulfilled by Lead Contact, Dr. David Talmage.

#### Materials Availability

Plasmids generated in this study have been deposited to Addgene and will be available upon publication under Talmage Lab.

#### Data and Code Availability

This study did not generate/analyze datasets. Code for fiber photometry data was previously published in (Crouse et al., 2020).

### EXPERIMENTAL MODEL AND SUBJECT DETAILS

Adult, male and female ChAT-IRES-Cre (B6;129S6-Chattm2(cre)Lowl/J, Jax stock number: 006410, (Rossi et al. 2011) mice were used for all DREADD experiments. In a subset of experiments used for ADCD labeling, these mice were crossed to Fos-tTA,Fos-EGFP* (TetTag, Jackson Laboratory; referred to as Fos-GFP). In all electrophysiology experiments, hemizygous Fos-tTA mice were used. Animals were housed in a 12-hour light/dark cycle environment that was both temperature and humidity controlled. Animals had free access to food and water. All animal care and experimental procedures were approved by the Institutional Animal Care and Use Committee of the SUNY Research Foundation at Stony Brook University and Yale University.

### METHOD DETAILS

#### Viral construct

##### Construction of the ADCD probe

All cloning unless otherwise specified was performed using In-Fusion HD (Clontech).

“mCherry-P2A” was amplified using Phusion High-Fidelity DNA Polymerase (NEB) from pV2SGE (obtained as a gift from Dr. Shaoyu Ge Stony Brook University). “oChIEF-LoxP-Lox2272” was amplified from pV2.2 (synthesized gene block from IDT). The two fragments were cloned into pAAV-WPRE linearized by BamHI. The resulting plasmid was linearized by Pml I. “7xTetO-LoxP-Lox2272-tTAH100Y.SV40” was amplified from pV2.1 (synthesized gene block from IDT) and cloned into the Pml I site. The final plasmid was packaged into AAV_9_ viral particles. Viral packaging was performed by the University of Pennsylvania Vector Core.

##### Construction of the ADCD-DREADD probe

“BglII-hM4Di.mCherry-AscI” was amplified using CloneAmpTM HiFi PCR Premix (Takara) from pAAV-hSyn-DIO-hM4D(Gi)-mCherry (Krashes MJ, et al. 2011) (gift from Dr.Bryan Roth; Addgene plasmid # 44362; http://n2t.net/addgene:44362; RRID:Addgene_44362). A backbone with TRE and Lox sites was ligated with “BglII-hM4Di.mCherry-AscI” using T4 DNA Ligase (NEB). The final plasmid was packaged into AAV_9_ viral particles. Viral packaging was performed by the University of North Carolina Vector Core.

##### Stereotaxic surgery & viral delivery

Three to four-month old ChAT-IRES-Cre mice were anesthetized and stereotaxically injected bilaterally. Coordinates were calculated based on the Paxinos Mouse Brain Atlas (Franklin.K & Paxinos.G, 1997): BLA (−1.4mm A/P, ±3.5mm M/L, −4.8mm D/V), NBM (−0.7mm A/P, ±1.7mm M/L, −4mm D/V).

###### Tracers

3% w/v solution of fast blue (FB) (17740-1, Polysciences Inc.) was prepared in sterile milliQ water. ∼0.2µL of 3% FB was injected into the BLA bilaterally of Fos-GFP mice. Mice were euthanized 7 days following injection.

#### Behavioral testing & analysis

##### Threat conditioning

All training and assessments were completed with experimenter blind to condition. Both training and recall sessions were analyzed using FreezeFrame v.3 (see below).

###### Habituation

All mice were handled for a minimum of five minutes daily for three consecutive days before behavioral training began. For DREADD experiments, all mice were additionally habituated to restraint and injection with 100 μL saline administered i.p. daily.

###### Training

On training day, all chambers were cleaned with 70% ethanol. Mice were placed into the behavioral chamber for a 10 min session which consisted of 3 min of habituation, followed by 3 tone-shock pairings (30 s 80dB, 5kHz tone, co-terminated with a 2 s 0.7mA foot shock with a 1.5 min interval between each pairing), and finally 2 min of exploration. For DREADD experiments, mice were given 0.1 mg/kg Clozapine (administered i.p.) (Sigma Aldrich) 10 minutes prior to being placed in the chamber.

###### Recall

Recall session took place 24 - 72 hrs after completion of the training. To specifically test the response to tone-cued recall, the contextual features of the chambers were altered including texture of the floor, color of the walls, and scent of cleaner (mild lemongrass citrus-based solution). Mice were placed in the behavioral chamber for another 5 min session during which a single tone was delivered (30 s 80dB 5kHz tone) 2 min after being placed in the chamber. No shock was administered.

###### Analysis

Percent time spent freezing was quantified using FreezeFrame v.3 (Actimetrics). Bout duration (defined as minimum required duration when animal is frozen) was set to 1 s, and threshold was defined as highest motion index with no movement other than breathing. Percent time spent freezing (defined as periods of no movement) was quantified across the 10 min session in bins of 30s. High responders were defined as those mice that exhibited at least a 10% increase in % time spent freezing in the bin during the tone from the bin immediately pre-tone. All other mice were considered low responders.

##### Engram labeling

Animals were placed on doxycycline hyclate-containing chow (Cat# TD.08541 Envigo) at least 2 days prior to injection of activity-dependent viral markers. Threat conditioning was performed as mentioned above. During doxycycline withdrawal, mice were transferred to a clean cage to prevent mice from eating dox food that was dragged into the cage or buried in the bedding. To minimize stress, some bedding containing fecal pellets and urine, and nest from the old cage were transferred to the new cage.

Following behavior during which labeling was desired, mice were returned to their old cages and left undisturbed for 72 hours.

##### Predator odor exposure

###### Habituation

All mice were habituated to restraint and injection with 100 μL saline administered i.p. daily for 3 days prior to behavioral testing for DREADD experiments. On exposure day, mice were transported to the lab several hours prior to exposure and habituated to the room and ambient sounds.

###### Exposure

For exposure to predator odors, a vented mouse cage (L 13in x W 7.5in x H 5.5in) with corncob bedding (EnviroDri) was placed in a designated location in a laminar flow hood with overhead fluorescent lighting. Mt. Lion Pee (Maine outdoor solutions LLC) was obtained from predatorpee.com and stored at 4°C. 200μL of urine was pipetted onto a 3in x 3in 12 ply gauze pad (Cat#6312, Dukal corp.) placed in a polystyrene petri dish (VWR) at the vented end of the cage. Mice were placed into the cage in the end away from the odor and the cage was covered using a plexiglass barrier. Mice were exposed for 5 minutes and the session was filmed using an overhead digital camcorder (Sony). Following exposure, mice were returned to their homecage or a holding cage in the case of multiple housed mice to prevent any odor transfer. Control mice were exposed to 0.9% saline. For DREADD experiments, mice were given 0.1 mg/kg clozapine (administered i.p.; Sigma Aldrich) 15 minutes prior to being placed in the chamber.

#### Fiber Photometry

##### Acquisition

Fiber photometry recordings were made using a Doric Lenses 1-site Fiber Photometry System. Signal was recorded using Doric Neuroscience Studio (V 5.3.3.4) via the Lock-In demodulation mode with sampling rate of 12.0 kS/s. Data was downsampled by a factor of 10 and saved as a comma-separated file. For details on connection of the setup refer to Crouse RB., et al. 2020.

##### Analysis

Preprocessing of the raw data was performed using a MATLAB script provided by Doric. The baseline fluorescence (F_0_) was calculated using a least mean squares regression over the duration of the recording session. The change in fluorescence for a given timepoint (ΔF) was calculated as the difference between it and F_0_, divided by F_0_, and multiplied by 100 to yield % ΔF/F_0_. The % ΔF/F_0_ was calculated independently for both the signal (465 nm) and reference (405 nm) channels and a final “corrected % ΔF/F_0_” was obtained by subtracting the reference % ΔF/F_0_ from the signal % ΔF/F_0_ at each timepoint. The corrected % ΔF/F_0_ was z-scored to give the final “Z % ΔF/F_0_” reported.

#### Electrophysiology

##### Brain slice preparation

For slice physiology, animals were anesthetized and transcardially perfused with cutting solution (sucrose 248 mM, KCl 2 mM, MgSO_4_ 3 mM, KH_2_PO_4_ 1.25 mM, NaHCO_3_ 26 mM, glucose 10 mM, sodium ascorbate 0.4 mM and sodium pyruvate 1 mM, bubbled with 95% O_2_ and 5% CO_2_) at 40°C. The brain was then rapidly removed and sliced, coronally, at 300 µM in oxygenated cutting solution at 40°C. Prior to recording, slices were incubated in oxygenated incubation solution (sucrose 110 mM, NaCl 60 mM, KCl 2.5 mM, MgCl_2_ 7 mM, NaH_2_PO_4_ 1.25 mM, NaHCO_3_ 25 mM, CaCl_2_ 0.5 mM, MgCl_2_ 2 mM, glucose 25 mM, sodium ascorbate 1.3 mM, and sodium pyruvate 0.6 mM) at room temperature.

##### Electrophysiological recording

During recording, slices were superfused with oxygenated artificial cerebral spinal fluid (Jiang et al. 2016). Fos+ neurons were identified by GFP expression. Signals were recording using patch electrodes between 4-6 MΩ, a MultiClamp 700B amplifier, and pClamp10 software. Pipette internal solution was as follows: 125 mM K-gluconate, 3 mM KCl, 1 mM MgCl_2_, 10 mM HEPES, 0.2 mM CaCl_2_, 0.1 mM EGTA, 2 mM MgATP, and 0.2 mM NaGTP (pH = 7.3). Following recording, cytoplasm was harvested via aspiration for cell-type identification using single-cell RT-PCR. Electrical properties were defined as previously described(López-Hernández et al., 2017).

#### Single cell reverse transcription-PCR

Single cell samples were pressure ejected into a fresh RT buffer prep (Applied biosystems). Samples were sonicated in a total volume of 20 μL at 40C for 10 min before addition of RT enzyme mix (Applied Biosystem). Tubes were incubated at 37^0^C for 60 minutes and then 95^0^C for 5 minutes. Two rounds of amplification (30 cycles each) were done for the detection of Chat transcripts. For the first round of amplification (reaction volume 25 μL) included 2X mastermix, sterile water, 0.2 μM of each primer, 1 μL of cDNA sample). For the second amplification, the reaction included 1 μL of the previous (first-round) PCR product, 2X mastermix, sterile water, and 0.2 μM of each primer. Whole brain cDNA was run in parallel with the single cell samples. After amplification, the PCR products (159 bp) were analyzed on 3% gels.

#### Immunohistochemistry

Following perfusion, brains were fixed overnight at 4°C in 4% PFA (in 1XPBS) and were then transferred to a 30% sucrose solution (in 1XPBS). Brains were flash frozen in OCT Compound (Tissue Tek) and stored at −80°C until cryosectioning. 50 µm cryosections were mounted onto Superfrost slides (Fisher Scientific) in sets of 3 and allowed to dry overnight at room temperature. Sections were blocked overnight at 4°C in a PBS solution containing 0.3% TritonX-100 and 3% normal donkey serum and then incubated with primary antibody in a PBS-T solution (0.1% TritonX-100 and 1% normal donkey serum), overnight (24h at 4 °C). The next day, sections were rinsed in PBS-T and incubated in secondary antibody for 2 hr at room temperature in PBS-T along with NeuroTrace-435 (Invitrogen). Sections were treated with an autofluorescence eliminator reagent (EMD Millipore) according to the manufacturer’s guidelines and mounted in Fluoromount-G (Southern Biotech).

### QUANTIFICATION AND STATISTICAL ANALYSIS

#### Imaging and analysis

All imaging used an Olympus VS-120 microscope at 20X magnification (Z-step= 3 um). Images were processed using the cell counter plugin on ImageJ. For Fos+ cell counts in the amygdala, only neurons (Nissl/ Neurotrace positive) with nuclear Fos stain were counted. The amygdala was identified and a region of interest (ROI) defined using ROI manager in Image J. Total area of the ROI was measured and noted. Fluorescence threshold was set to eliminate background fluorescence in ImageJ (defined as hazy background signal detected in space between neurons and white matter). This eliminated non-specific fluorescence and out of focus signals. Fos+ nuclei were then counted using the cell counter plugin.

For ADCD cell counts, neurons with ADCD label from the NBM injection site were counted. NBM was identified as the cluster of cholinergic cell bodies at the base of the internal capsule in the Globus Pallidus as denoted by the Paxinos Mouse Brain Atlas (3^rd^ Edition). 100% of the analyzed area of every third brain section was counted (∼150 um apart).

#### Statistical analysis

Statistical analysis was done using GraphPad Prism (GraphPad Software Inc., San Diego, CA, USA), Sigmaplot 12.5 (Systat Software, Inc., San Jose, CA, USA) and OriginPro 9.1 (Origin Lab Corporation, Northampton, MA, USA). Normality of the data was assessed using Shapiro-Wilk and Smirnov-Kolmogorov tests. Data that failed either normality test were analyzed using appropriate non-parametric tests. Detailed information on statistical tests used, p-values, and sample sizes can be found in the text (Figure Legends). Power was >0.9 for all reported data.

## Supplemental Figures

**Figure S1 (Related to Figure 1)**

**A.** ACh release in response to 3 sequential 30 second tones (without shock; n=5). Quantification of change in slope (left) and maximum ACh release (right), comparing the baseline period for each of the three tones. There was no significant change in ACh release (slope or peak) in response to any of the three tones. Friedman test p=0.0933 (slope) p=0.2982 (peak).

**B.** ACh release in response to tone 24 hrs following exposure to three tones without foot shock (tone without shock) (n=5). Quantification of change in slope (left) and maximum ACh release (right), comparing the baseline period to naïve tone (tone 1), and “recall” tone. There was no change in ACh release in response to the “recall” tone following training with tone alone (no foot shock). Friedman test p=0.1092 (slope) p=0.3673 (peak).

**Figure S2 (Related to Figures 2&3)**

**A. Diagrams of ADCD and ADCD hM4Di viral constructs.** Activity dependence is conferred by the <u>T</u>et response <u>e</u>lement (TRE – 7 repeats of the tetO sequence followed by a minimal promoter). Cre-dependence is conferred by pairs of loxP and lox2272 sites flanking the “cargo” in an antisense orientation. Cargos: permanent labeling ADCD includes an oChIEF-mCherry fusion followed by a P2A element and a doxycycline insensitive tTA (tTA*). ADCD-hM4Di cargo includes a hM4Di-mCherry fusion protein.

**B. Test to determine the minimal time off dox to allow ADCD expression.** Animals were kept on dox food for 2 weeks (starting 2 days prior to ADCD+AAV-Cre-IRES-GFP injection into the BLA) (Dox ON). They were then shifted to regular chow either 6, 12 or 24 hr prior to training (Dox OFF). They were returned to dox chow immediately following training (ON). ADCD expression was quantified 72 hr later as % of mCherry+ cells out of total GFP/Cre+ cells in the BLA.

**C.** Images showing lack of ADCD expression in Chat-Cre and Fos-tTA mice, and ADCD expression in Chat-Cre X Fos-tTA mice. Scale bar 50 µm.

**Figure S3 (Related to Figure 4).**

**A.** Quantification of BLA-projecting cholinergic neurons (BFCNs) labeled by injection of CAV_2_-DIO-hM4Di.mCherry into the BLA of Chat-IRES-Cre mice along the A-P axis (black boxes, left Y axis) and of recall-activated cholinergic neurons labeled using ADCD virus as mentioned in Fig 4 along the A-P axis (red boxes, right Y axis).

**Figure S4 (Related to Figure 5).**

**A. Left**. Strategy for retrograde targeting of hM4Di DREADD to BLA-projecting cholinergic neurons. BLA-projecting cholinergic neurons were silenced by injection animals with clozapine (CLZ) prior to training.

**B.** Inhibiting BLA projecting cholinergic neurons during training preferentially prevented recall activation of BLA neurons (Fos+ density) in anterior regions of the BLA, between Bregma −0.8 and −1.4. **Black dots** – control v. **red dots** – cav.hM4Di^BLA^

**C.** Silencing BLA-projecting cholinergic neurons during **RECALL** reduced CeC Fos immunoreactivity profiled following recall. Images of CeC Fos immunoreactivity in sham control vs. CAV_2_-DIO-hM4Di.mcherry (image taken at A/P = −0.8 from Bregma). Fos+ cell density in CeC of control vs. CAV_2_-DIO-hM4Di.mcherry injected animals (n=3 mice/group, 6 sections control vs. 7 sections hM4Di). Mann-Whitney test *: p=0.0012.

**D.** BLA-projecting cholinergic neurons were silenced by injecting animals with clozapine (CLZ) during training. Percent time freezing during the recall session including the pre-tone period (average over 90 sec), during the tone (30 s) and during the 90 s post-tone period. Clozapine was only administered during the training session. RM two-way ANOVA (GLM) w/ post-hoc Bonferroni correction. * denotes a significant difference between tone and post-tone vs. pre-tone for controls (p<0.05). # denotes significant difference between groups over recall session (p<0.05).

**E.** BLA-projecting cholinergic neurons were silenced during recall (clozapine given **ONLY** during the recall. Freezing differed significantly between pre-tone v. tone and post-tone period for controls (RM two-way ANOVA (GLM) w/ post-hoc Bonferroni correction *, p<0.05). # denotes significant difference between groups over recall session (p<0.05).

**F.** Clozapine was injected in control animals, (black, no hM4Di) and animals expressing hM4Di in BLA-projecting cholinergic neurons (red) immediately following the training session. Freezing behavior was measured 24 hrs later. There was no difference in freezing during pre-tone, tone or post-tone periods between control animals and animals in which BLA-projecting cholinergic neurons has been silenced during early consolidation. Control: n=3 mice, hM4Di: n=4 mice.

**G.** Silencing BLA-projecting cholinergic neurons increases the proportion of low responding animals. Control and hM4Di expressing mice to which clozapine was injected during training (left) or during recall (right) were stratified into low (black fill) and high (white fill) responders. Silencing BLA-projecting cholinergic neurons during training resulted in 100% low responders during recall (control 1 of 5 low responders; hM4Di 7 of 7 low responders), whereas silencing this population during recall shifted the proportion of low responders from 10% in the control (1 of 10) to 50% (3 of 6).

**H.** Silencing CeA-projecting cholinergic neurons during recall had no effect on freezing behavior. (left) hM4Di DREADD was targeted to CeA-projecting cholinergic neurons (red, controls in black); and these neurons were silenced during recall. No significant differences were found between groups (Control: n=9; hM4Di: n=4).

**Figure S5 (Related to Figure 6)**

**A-F:** Population data presented as dot plot + median for the electrophysiological properties of post hoc identified cholinergic neurons from home cage (HC, n= 11 ChAT+ neurons from 11 mice) compared with those of Fos- and Fos+ NBM cholinergic neurons following recall to tone alone (Fos-: n= 10 ChAT+ neurons from 5 mice; Fos+: n= 11 ChAT+ neurons from 6 mice,). Kruskal-Wallis tests **A-F**: action potential amplitude, p>0.05 (**A**); threshold, p>0.05 (**B**); latency, p = 0.0004 (Dunn’s Corrected p-values: HC vs. Fos-: p = 0.0009, HC vs. Fos+: p = 0.0004, Fos- vs. Fos+: p = 0.7394) (**C**); resting membrane potential, p>0.05 (RMP, **D**); after hyperpolarization (AHP) amplitude, p = 0.0174 (Dunn’s Corrected p-values: HC vs. Fos-: p = 0.3702, HC vs. Fos+: p = 0.0041, Fos- vs. Fos+: p = 0.0952) **E**; and AHP half-width, p > 0.05 (**F**).

**G:** Population data presented as dot plot + median for the maximum firing rate of post-hoc identified cholinergic neurons from home cage (n=11 ChAT+ neurons from 11 mice) and immediately following acquisition (n=5 ChAT+ neurons from 3 mice).

**H:** Population data presented at dot plot + median for max firing rate of post hoc identified cholinergic neurons from home cage, training, 3h post-recall (D0), and ADCD+ NBM neurons identified 3 days post-recall (D3) and 5 days post-recall (D5) from Chat-IRES-Cre x Fos-tTA mice injected with ADCD-DIO-oChIEF.mCherry with Dox off during recall. Kruskal-Wallis tests, p = 0.0072 (D0 vs. all other groups p<0.01)

**Figure S6 (Related to Figure 7).**

**A.** Number of contacts with odor pad (left) and number of digging bouts (right) during a 5 min saline or predator odor exposure (saline, gray: n=9 mice and predator odor, black: n=6 mice). Animals displayed defensive behaviors, including avoiding the odor pad and increased digging bouts in response to the predator odor. Mann-Whitney test ***: p= 0.002; *: p= 0.026.

**B.** Number of contacts with odor pad (left) and digging bouts (right) during a 5 min predator odor exposure in clozapine injected animals (control n=6 mice and hM4Di n=5 mice). Silencing BLA-projecting cholinergic neurons did not affect avoidance behavior (# pad contacts) or digging.

**C.** The NBM/SI_p_ of Chat-Cre mice was directly targeted with an AAV-hM4Di DREADD and cholinergic neurons were silenced during recall. Freezing behavior during recall (pre-tone period, tone and post-tone). Controls (black) froze in response to tone (RM two-way ANOVA (GLM) w/ post-hoc Bonferroni correction **, p<0.01). Silencing the NBM/SI_p_ significantly reduced freezing across the session (control vs hM4Di: #, p<0.05. Control, n=5, hM4Di n=4).

**D.** hM4Di DREADD was targeted to VP/SI_a_ cholinergic neurons; and clozapine was administered during the recall session. Silencing VP/SI_a_ cholinergic neurons during recall had no effect on freezing behavior. Control: n=3, hM4Di: n=4.

## Notes

### Competing Interest Statement

The authors have declared no competing interest.

### Summary of Updates

Figures resized to accommodate copyright text.

