## Supplemental Figures for "Basal forebrain cholinergic neurons are part of the threat memory engram"

### Supplemental Figures S1-S6

### Figure S1 (Related to Figure 1)

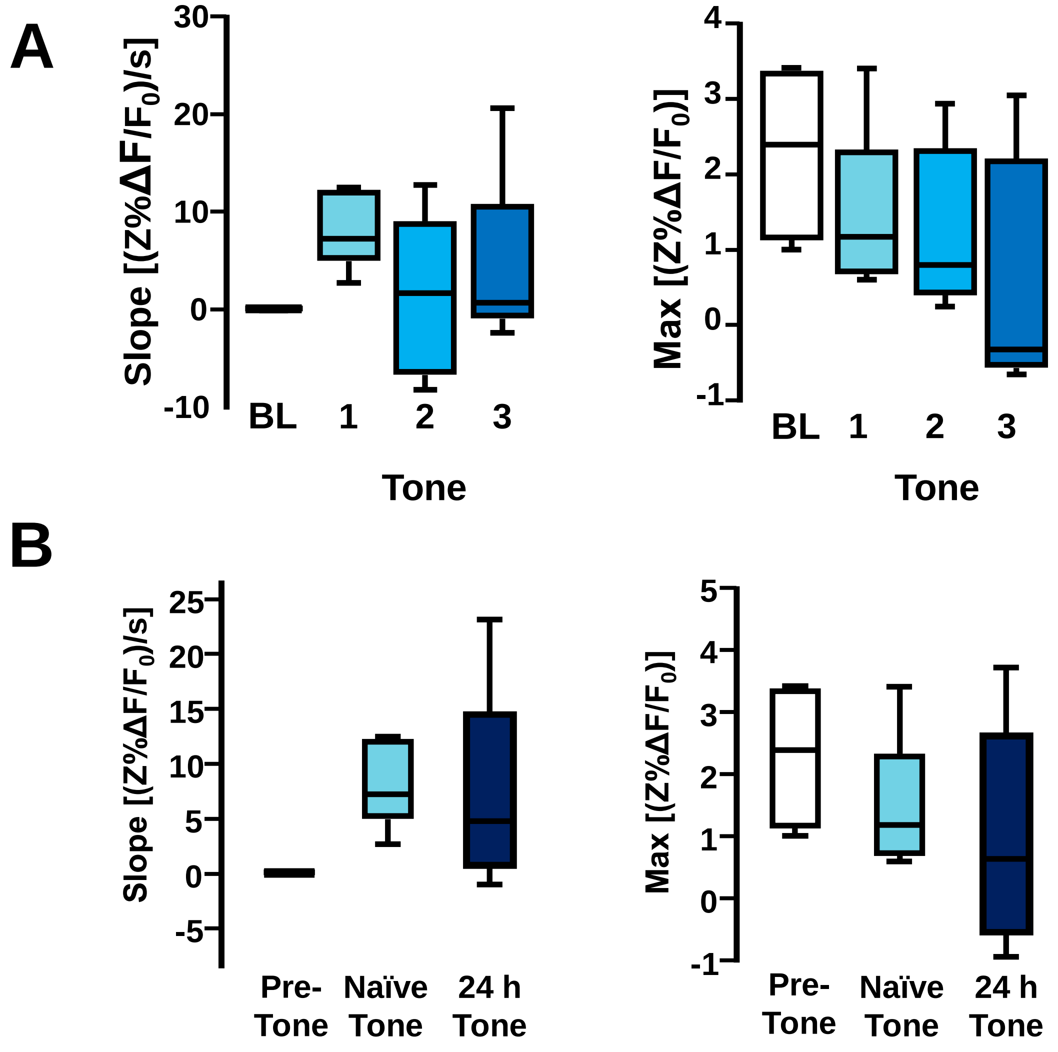

### Figure S2 (Related to Figures 2 and 3)

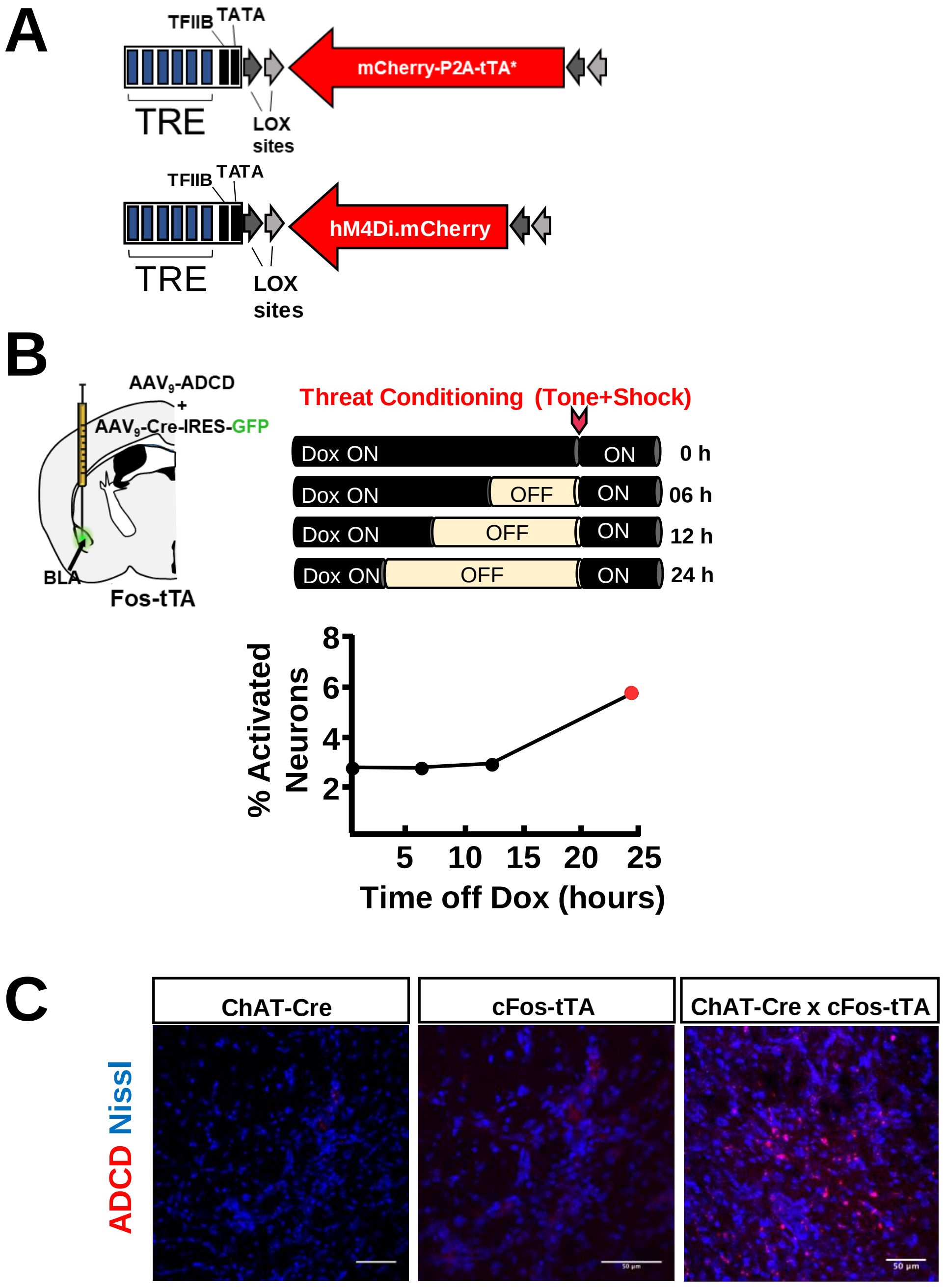

### Figure S3 (Related to Figure 4)

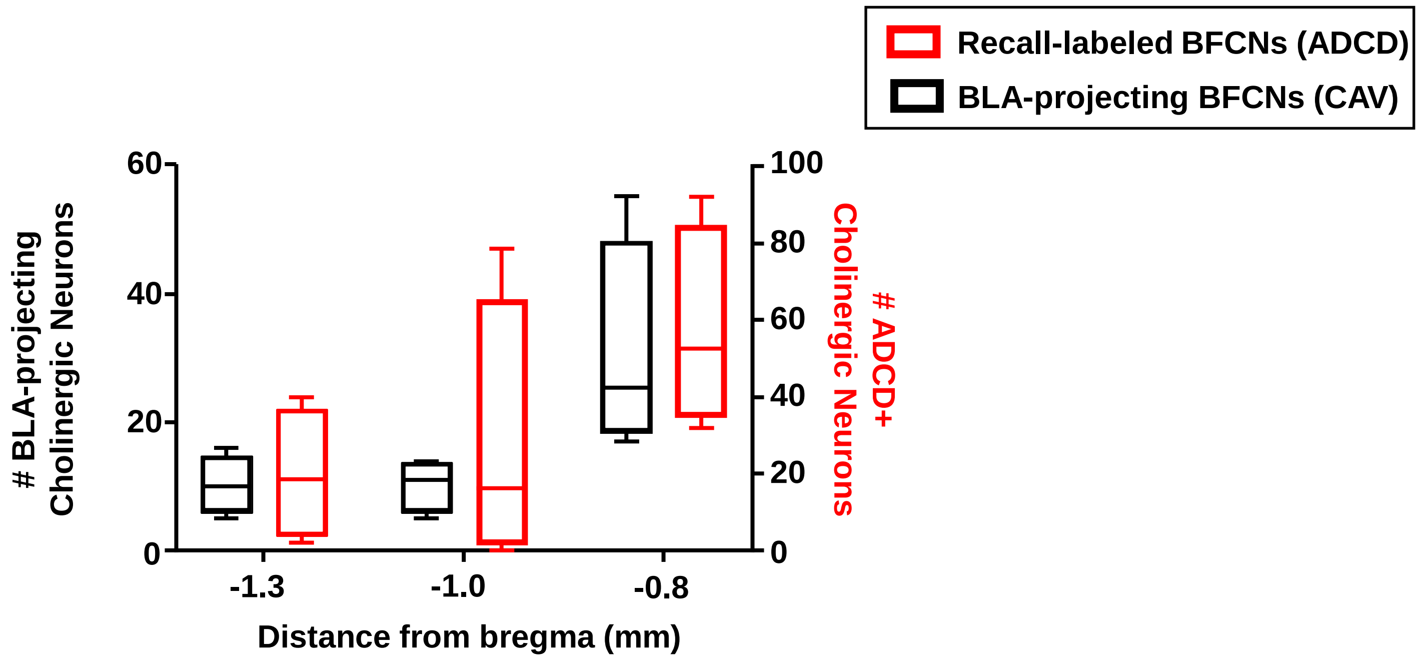

### Figure S4 (Related to Figure 5)

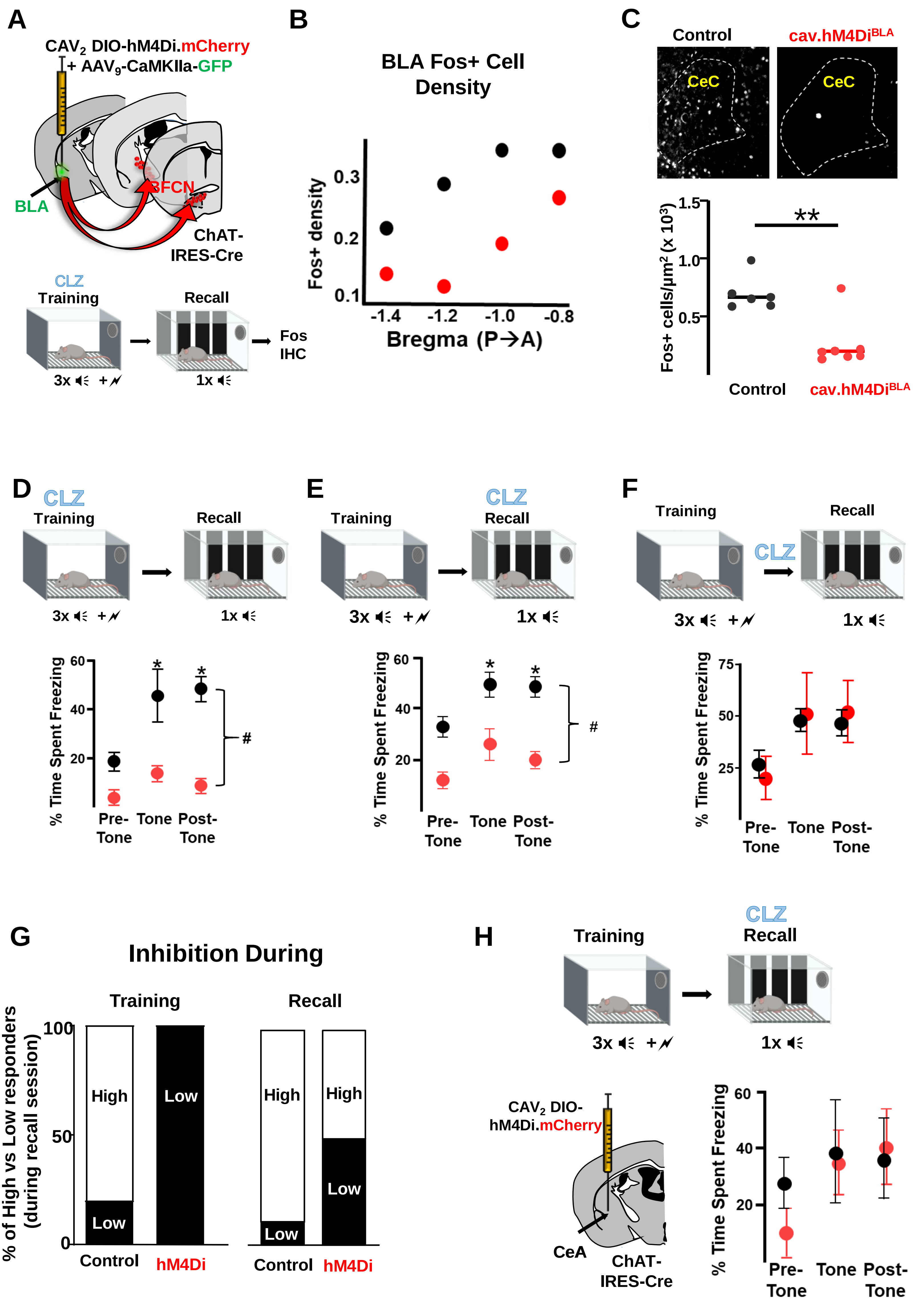

### Figure S5 (Related to Figure 6)

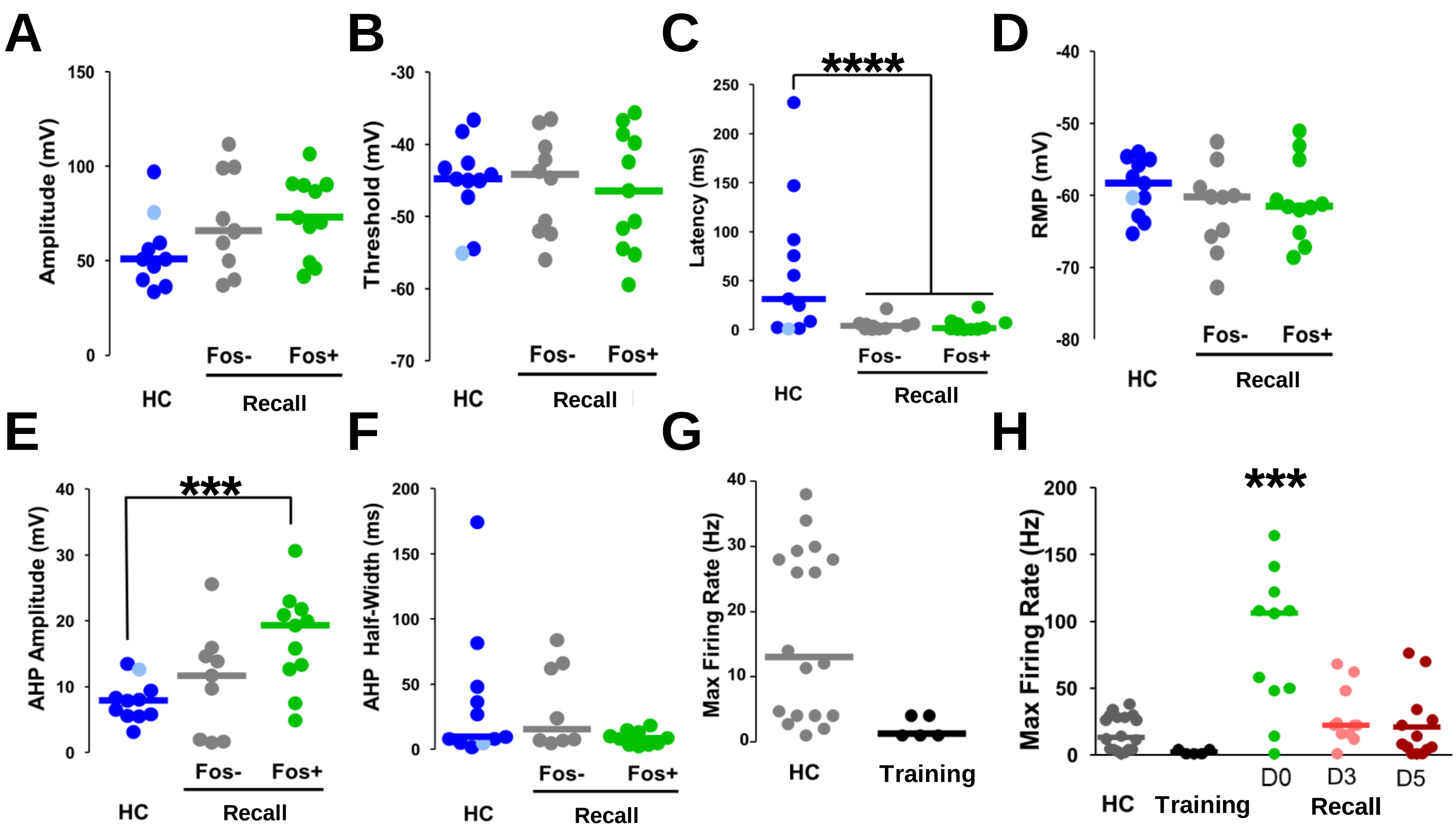

### Figure S6 (Related to Figure 7)

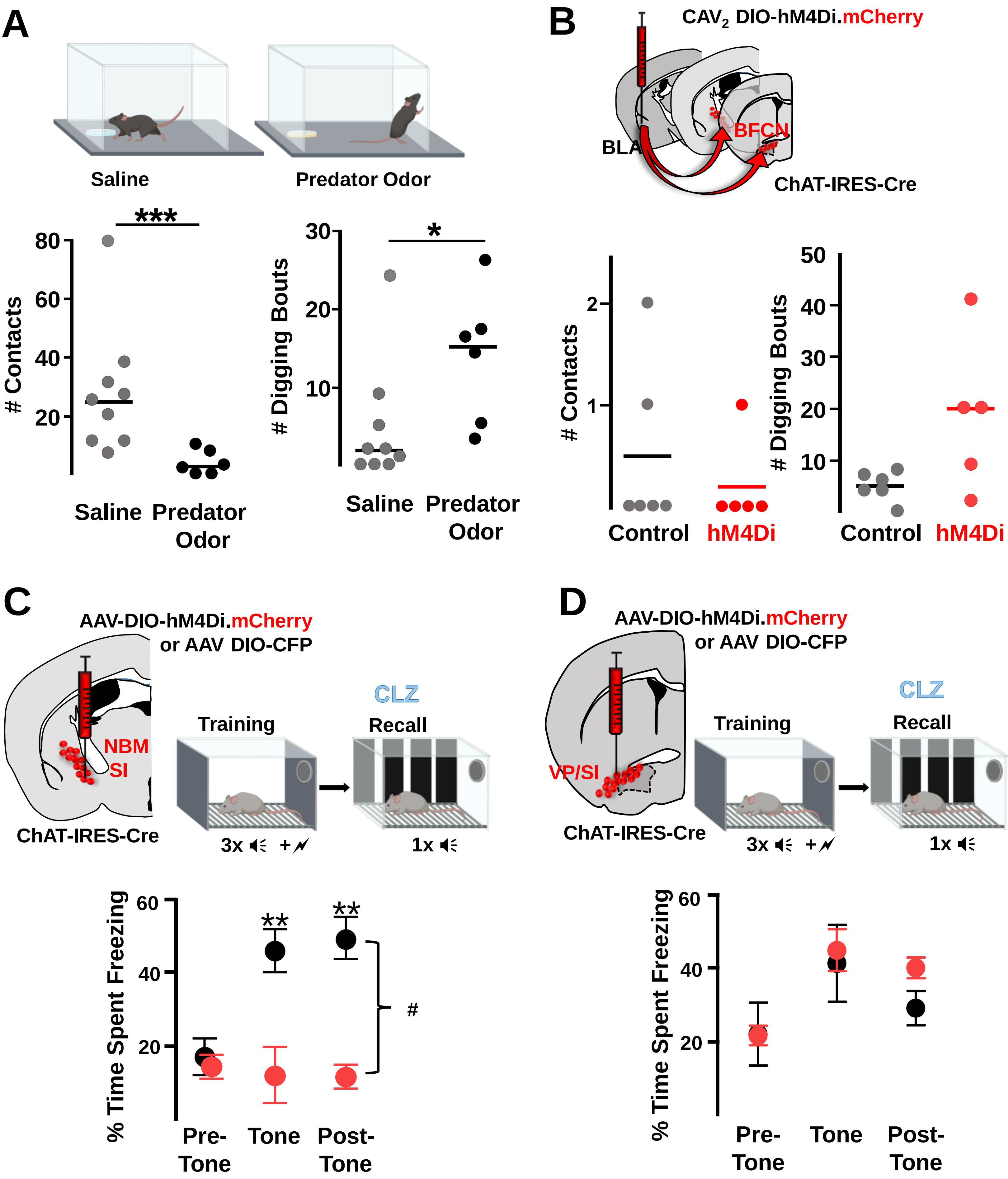
